# Precision and Accuracy in Quantitative Measurement of Gene Expression from Single-Cell/Nuclei RNA Sequencing Data

**DOI:** 10.1101/2024.04.12.589216

**Authors:** Rujia Dai, Ming Zhang, Tianyao Chu, Richard Kopp, Chunling Zhang, Kefu Liu, Yue Wang, Xusheng Wang, Chao Chen, Chunyu Liu

## Abstract

Single-cell and single-nucleus RNA sequencing (sc/snRNA-Seq) have become essential tools for profiling gene expression across different cell types in biomedical research. While factors like RNA integrity, cell count, and sequencing depth are known to influence data quality, quantitative benchmarks and actionable guidelines are lacking. This gap contributes to variability in study designs and inconsistencies in downstream analyses. In this study, we systematically evaluated quantitative precision and accuracy in expression measures across 23 sc/snRNA-Seq datasets comprising 3,682,576 cells from 339 samples. Precision was assessed using technical replicates based on pseudo-bulks created from subsampling. Accuracy was evaluated using sample-matched scRNA-Seq and pooled-cell RNA-Seq data of mononuclear phagocytes from four species. Our results show that precision and accuracy are generally low at the single-cell level, with reproducibility being strongly influenced by cell count and RNA quality. We establish data-driven thresholds for optimizing study design, recommending at least 500 cells per cell type per individual to achieve reliable quantification. Furthermore, we show that signal-to-noise ratio is a key metric for identifying reproducible differentially expressed genes. To support future research, we developed VICE (Variability In single-Cell gene Expressions), a tool that evaluates sc/snRNA-seq data quality and estimates the true positive rate of differential expression results based on sample size, observed noise levels, and expected effect size. These findings provide practical, evidence-based guidelines to enhance the reliability and reproducibility of sc/snRNA-seq studies.

## Introduction

Single-cell/nuclei RNA sequencing (Sc/snRNA-Seq) is a powerful technology developed for measuring gene expression in individual cells. The first scRNA-Seq study was published in 2009 by Tang et al[1]. Smart-seq was developed, enabling the amplification and sequencing of full-length mRNA transcripts from individual cells, characterizing transcriptomes at single-cell resolution. Since then, more technologies have been developed for single-cell profiling[2], with 10X chromium and Smart-seq being the two most commonly used methods.

Sc/snRNA-Seq has been used in various applications, including identifying novel transcriptional regulatory mechanisms[3], characterizing cell types and tissue compositions[4], studying developmental dynamics and trajectory of different cell types[5, 6], and identifying cell-type-specific changes as biomarkers for disease or treatment responses[7–9]. All these studies rely on accurate and precise measures of gene expression in each cell type. Precision and accuracy in the quantitative measurement of gene expression are defined as the variability of expression across replicates and the degree to which expression measurements match the actual or true values, hereafter referred to as ‘precision’ and ‘accuracy,’ respectively. Only when gene expression is quantified precisely and accurately in each sample, can the results of downstream analyses be reproducible and meaningful.

Random and systematic technical variability adds noise to the expression measurements in sc/snRNA-Seq[10]. Many zero values are observed in sc/snRNA-Seq data, called ‘dropouts’[11]. Dropouts can be caused by target genes truly not being expressed, or due to technical factors such as low mRNA input, mRNA degradation, capture efficiency, amplification efficiency, and sequencing depth. These technical factors can reduce precision and cause bias in accuracy of gene expression measurements. Previous studies attempted to assess technical noise in scRNA-Seq data using Spike-ins[12], sample-matched bulk-tissue RNA-Seq data[13], or qPCR[14] as references. However, these methods have rarely been used due to costs and practical limitations.

Strategies to improve the quality of single-cell data such as pooling more cells have been developed, but standardized procedures for completing sc/snRNA-Seq are lacking. There is a lack of systematic, quantitative thresholds to guide experimental design, making it challenging to define optimal parameters for achieving reliable results. These factors are often inconsistently evaluated across published studies, resulting in variability in data quality assessment. Practical guidelines—such as the minimum number of cells required per cell type—are either lacking or too vague, leaving researchers without clear direction for ensuring robust data quality in their experiments.

We evaluated the precision and accuracy of expression measurements with 23 sc/snRNA-Seq datasets produced on three different platforms published in high-profile journals (Table 1) in the framework as illustrated in **Figure 1**. Initially, we surveyed the cell numbers and missing rates in these sc/snRNA-Seq data, followed by calculating precision in each dataset using technical replicates based on pseudo-bulks. Additionally, we explored the impact of several technical factors, including RNA quality, saturation rate, total reads, and sequencing platform on expression precision. We also evaluated the expression accuracy with four datasets of cultured mononuclear phagocytes from sample-matched pooled-cell RNA-Seq and scRNA-Seq data.

**Figure 1.**
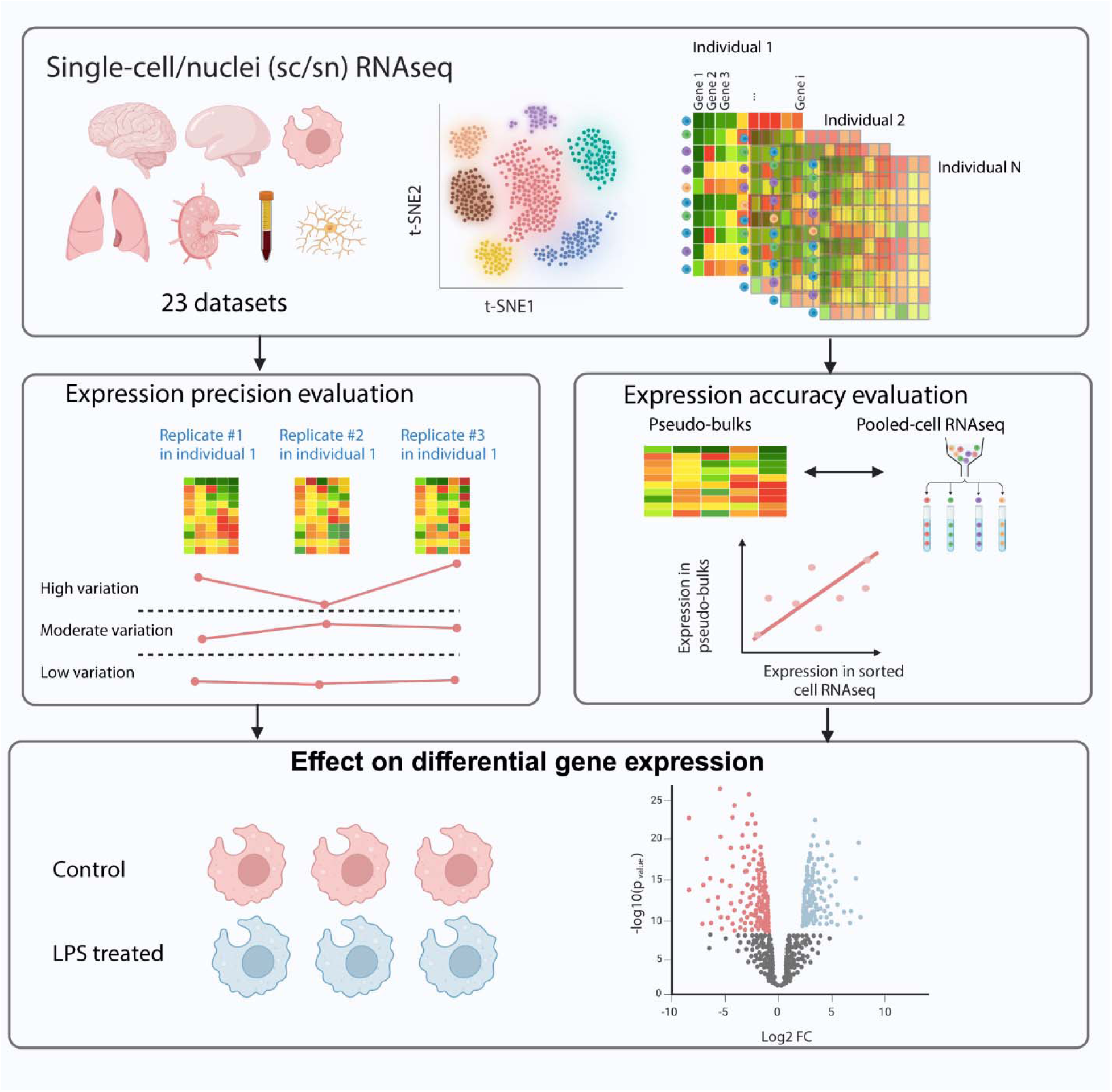
Overview of the study. Framework for evaluating the expression precision and accuracy of sc/snRNA-Seq across datasets and platforms. We assessed the precision and accuracy of gene expression measurements using 23 sc/snRNA-Seq datasets generated on three platforms. These datasets, published in high-profile journals, were derived from large consortium efforts, including the BICCN, reflecting current technological standards. Our analysis began with a survey of cell numbers and missing rates across datasets, followed by the evaluation of precision based on technical replicates. We then examined the influence of technical factors such as RNA quality, sequencing saturation rate, total read counts, and platform type on expression precision. To assess accuracy, we compared scRNA-Seq data from four cultured mononuclear phagocyte datasets with corresponding pooled-cell RNA-Seq data from the same samples. Finally, we analyzed the effects of cell numbers and other factors on the reproducibility of downstream differential expression analyses. This figure was created with BioRender.com. Sc/snRNA-Seq, Single-cell/nuclei RNA sequencing expression measures; BICCN, BRAIN Initiative Cell Census Network.

**Table 1.**
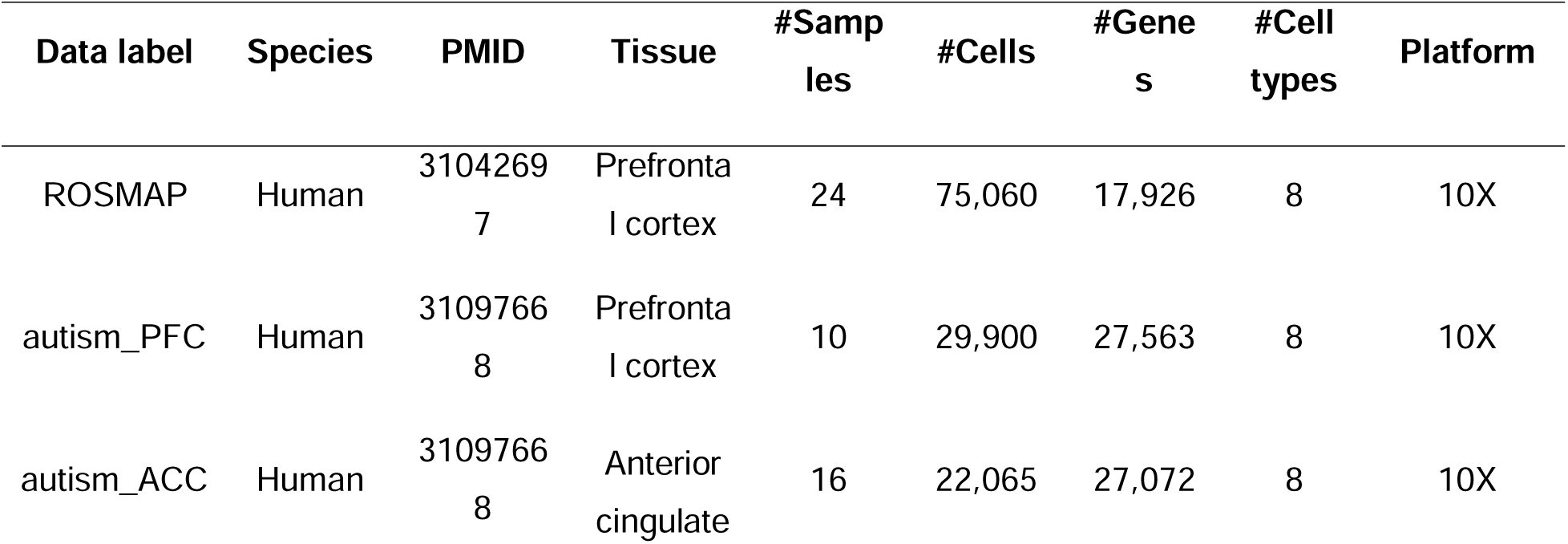
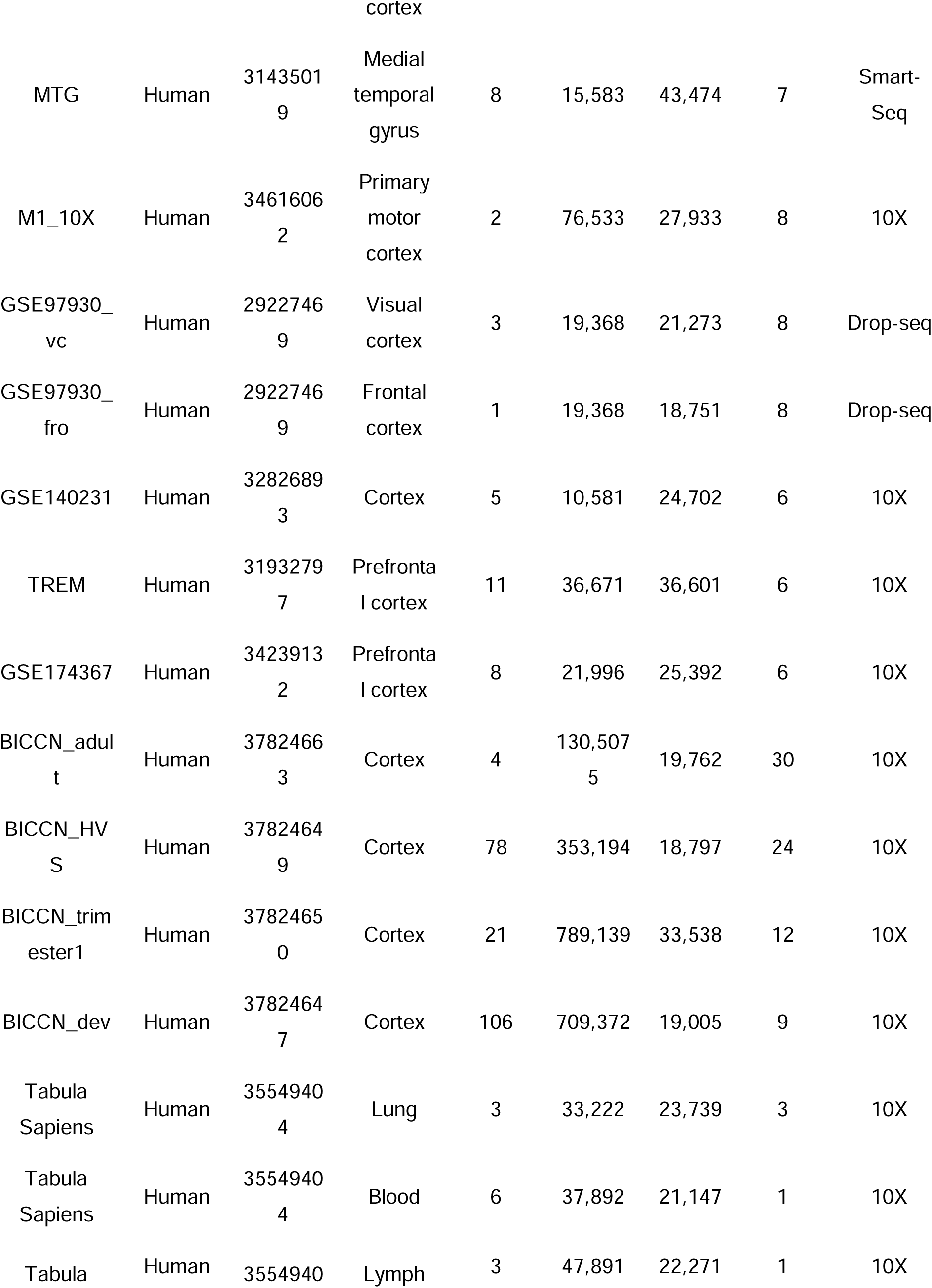
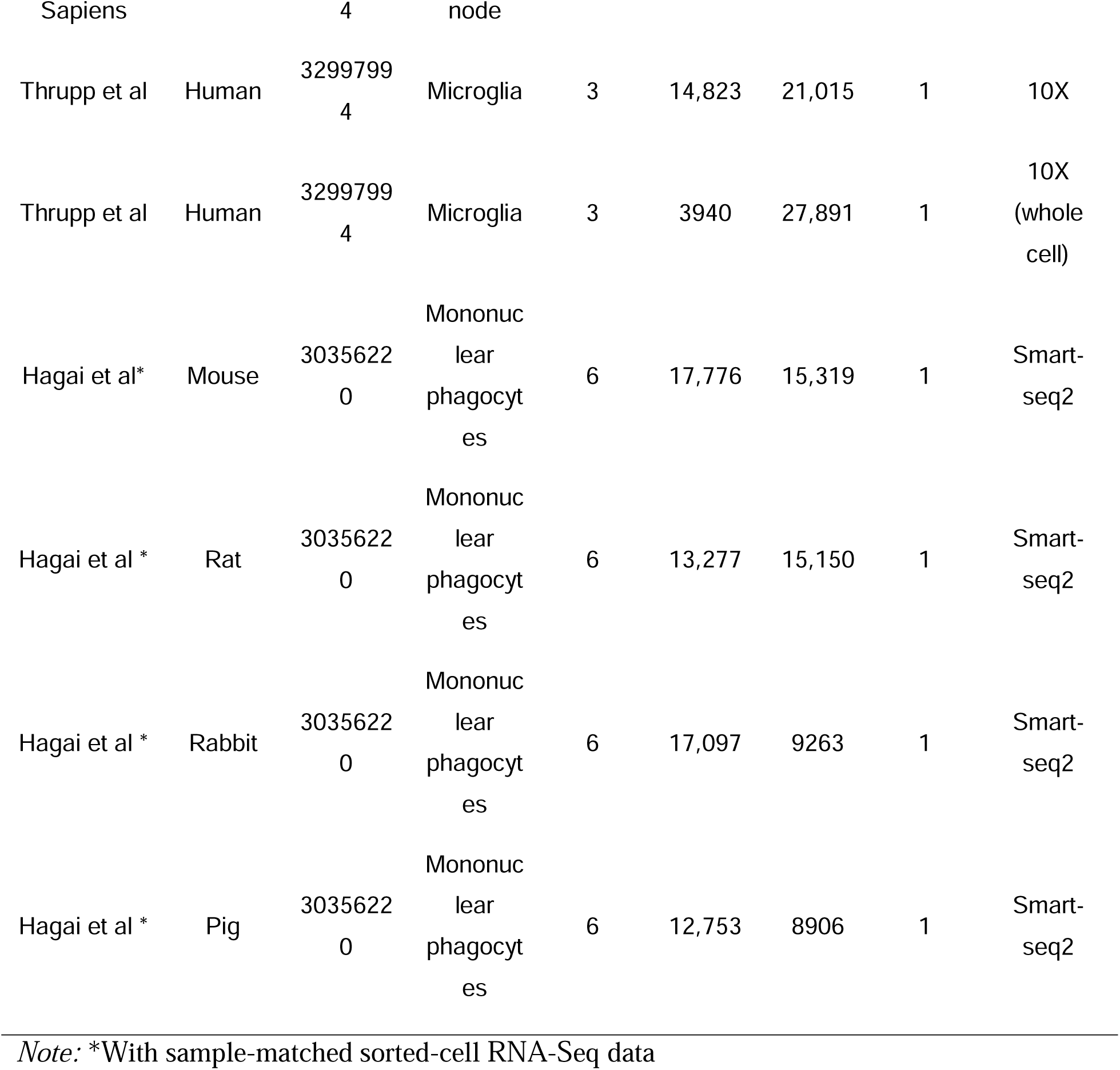
Single-cell/nuclei RNA sequencing datasets assessed in this study.

Lastly, we evaluated the effect of cell number and other factors on the reproducibility of downstream differential expression (DE) analysis. Based on the evaluation, we provided practical guidelines for future studies. To facilitate future experiment design and data evaluation, we developed a tool we named VICE (Variability In single-Cell gene Expressions) https://github.com/RujiaDai/VICE.

## Results

### Existing sc/snRNA-Seq data have high missing rate

We measured the missing rate for each gene at both the individual-cell and pseudo-bulk levels. Pseudo-bulks were created from single-cell gene expression of a specific cell type within an individual to mimic bulk RNA-Seq data. The missing rate was defined as the proportion of cells with zero expression for a given gene across all individual cells or pseudo-bulks of the same cell type. Individual cells had an average missing rate of 90% (**Figure 2A**), while the pseudo-bulks reduced the average missing rate to 40% (**Figure 2B**). Including more cells in the pseudo-bulks resulted in a lower observed missing rate (Figure S1).

**Figure 2.**
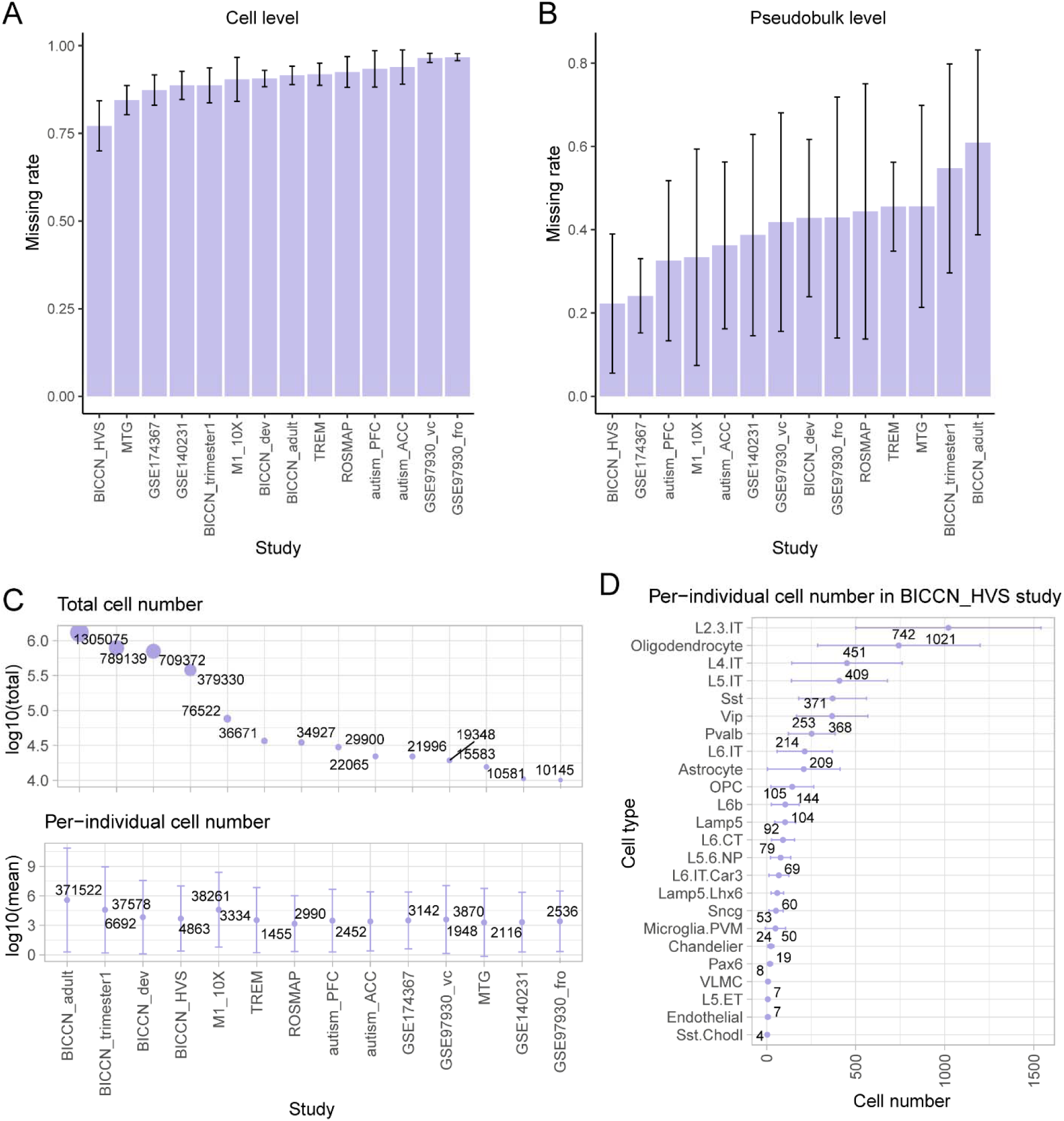
Missing rates and cell numbers in the 14 datasets. **A**. Missing rate per gene at individual cell level. **B**. Missing rate per gene at the pseudo-bulk level. **C**. Total number of cells studied, along with the average number of cells per individual. The cell count per individual was calculated by dividing the total number of cells by the number of individuals in the study. Cell numbers were log10-transformed for better visualization. D. An illustrative example showcasing the average number of cells per individual, specifically drawn from the BICCN_HVS study.

Though each project sequenced many cells, we noticed that the number of cells sequenced per cell type per individual was sometimes very small, particularly for minor cell types. Across the 14 brain datasets, the average total cell count was 247,190, whereas the average cell number per individual was 34,483 (**Figure 2C**). The number was even much smaller for specific cell types per individual. For instance, the BICCN_HVS study sequenced 353,194 cells and categorized 24 cell classes (**Figure 2D**). The largest group of cells in this data comprised an average of 1,021 intratelencephalic (IT) neurons from layers 2 and 3, while the smallest group had only an average of 4 somatostatin (SST) chodl inhibitory neurons across the samples, a difference of three orders of magnitude.

### Low expression precision in sc/snRNA-Seq data

Expression precision was evaluated by the expression variability across technical replicates based on pseudo-bulks in sc/snRNA-Seq data. First, we generated technical replicates based on pseudo-bulks by randomly grouping cells of the same type from the same individual into three groups and totaling expression values of each gene from all cells within each group (Figure S2). We then calculated the coefficient of variation (CV) for each gene to measure the variability of gene expression across the technical replicates based on pseudo-bulks in each cell type. To avoid sampling bias, we calculated the CV 100 times and used the averaged CV to represent the overall precision in the data.

Our analysis revealed that the median CV of detected genes across technical replicates based on pseudo-bulks was 0.68 ± 0.24 in the 14 brain datasets (**Figure 3**), which is much higher than the median CV observed in bulk-tissue RNA-Seq[15] (ranging from 0.11 to 0.39) and microarray data[16] (ranging from 0.1 to 0.2). Utilizing cell classification in the BICCN_HVS study, we calculated the CV values at both cell type and subtype levels. The CVs were not significantly different at these two resolution levels, suggesting the observed variability was not driven by heterogeneity in a higher-level cell classification (Figure S3). A similar pattern was noted in independent mouse brain data when different numbers of cells were sequenced, indicating that low precision is a technical challenge in single-cell data across various sample sources (Figure S4).

**Figure 3.**
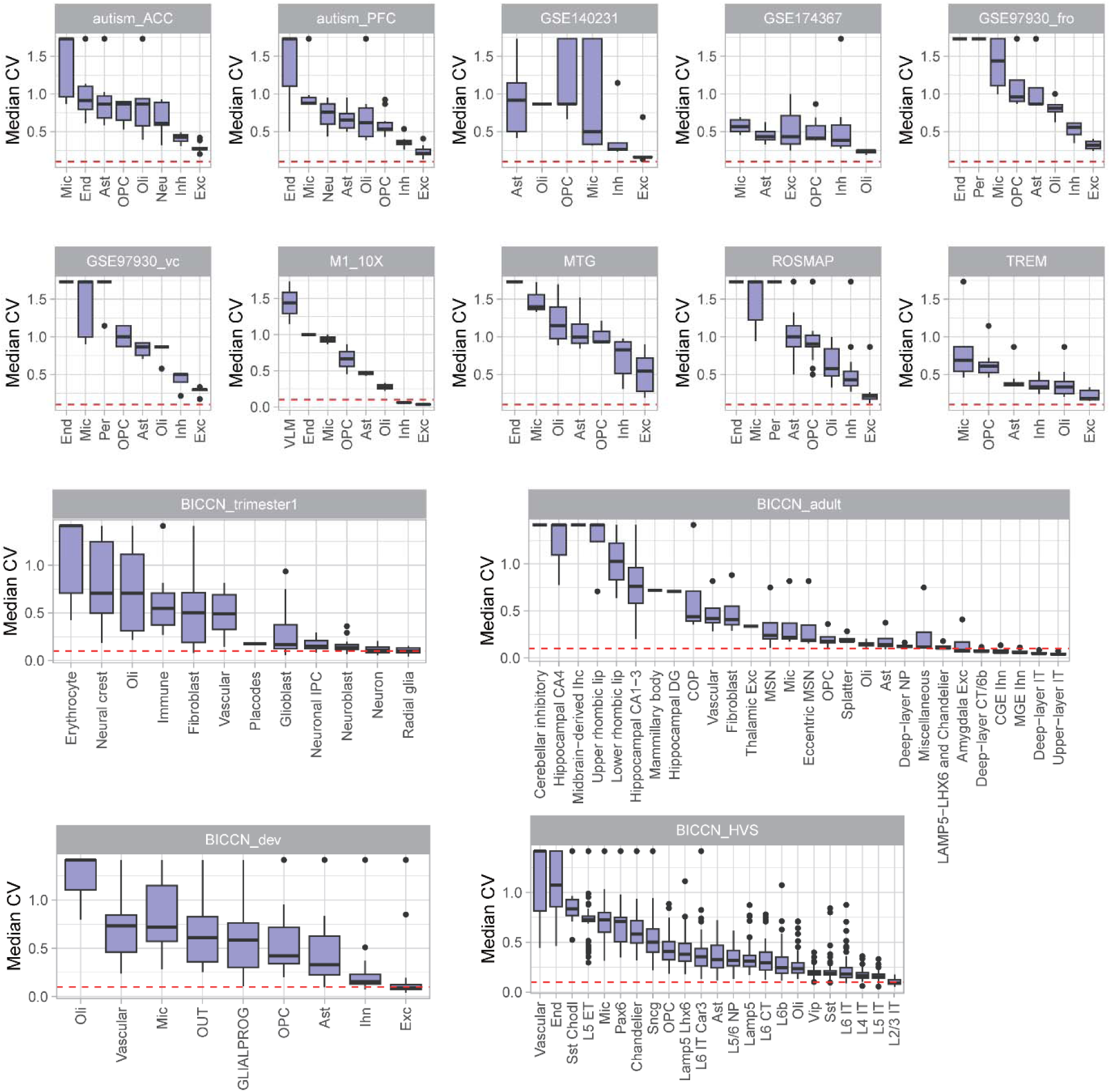
Gene expression precision evaluated by technical replicates in sc/snRNA-Seq data. We calculated gene expression variability as the CV of gene expression in three technical replicates based on pseudo-bulks for each gene. The sample with the largest number of total cells in each dataset was used for illustration. The red dashed line denotes the CV thresholds of 0.1, which is a threshold recommended for bulk-tissue gene expression quality control processing[16]. CV, coefficient of variation.

To illustrate the expression variability in multiple tissues, the single-cell RNA-seq data from blood, lung, and lymph nodes were evaluated[17]. A similar CV pattern across cell numbers was observed, consistent with findings in brain tissue data. Regardless of tissue type and cell type, approximately 500 cells are needed to drive CV close to 0.1 (Figure S5).

Major cell types exhibited lower CV than minor cell types. For example, excitatory neurons, as the most abundant cell type, had a CV of 0.19±0.20 across datasets. In contrast, the other cell types had a median CV of 0.55±0.40 across datasets, indicating that the precision problem is particularly severe for low-abundance cell types. Additionally, expression CV was negatively correlated with expression abundance (Correlation coefficient = −0.88, p-value < 2.2e-16, Figure S6A). Notably, marker genes have lower CV than other genes (Figure S6B).

To compare the expression variability in sc- and snRNA-Seq data, we evaluated three brain microglia samples with both sc- and snRNA-Seq data[18]. We observed almost identical CV patterns in the two data types, indicating that quality issues are a common concern for both (Figure S7). We calculated the percentage of samples achieving a designated precision threshold, a CV of 0.1 or lower for each cell type. There was a striking disparity: the proportion of samples from five distinct datasets that satisfied this precision criterion ranged from 3% to 25%, with an average of 5%, as illustrated in Figure S8A. For example, every sample representing upper-layer IT neurons in the BICCN adult dataset successfully passed the precision assessment (Figure S8B). In the case of the BICCN_HVS dataset, 67% of samples pertaining to IT neurons in layers 2 and 3 met the established quality benchmarks (Figure S8C). Conversely, in the other nine datasets, not a single sample reached the requisite levels of precision. This indicates a prevalent problem with gene expression noise in individual samples of these datasets.

### Expression precision is correlated with number of cells sequenced

We expected that the expression precision would be associated with the number of cells sequenced and aimed to identify the minimum cell number for acceptable precision. To prove the expectation by actual data, we generated technical replicates based on pseudo-bulks with varying numbers of cells, ranging from single cell to the maximum cell number divided by three. The sample with the largest number of total cells in each dataset was utilized for testing. As the number of cells pooled into the technical replicates based on pseudo-bulks increased, the overall variability decreased until it reached a plateau for major cell types (**Figure 4A** and **Figure 4C**). With the small total number of cells sequenced, the minor cell types did not reach a stable CV (Figure S9). Similar correlation coefficients of −0.66±0.07 and −0.78±0.14 were observed between the number of cells in each replicate and median CV in excitatory neuron and oligodendrocyte respectively (p-value < 0.05, **Figure 4B** and **Figure 4D**).

**Figure 4.**
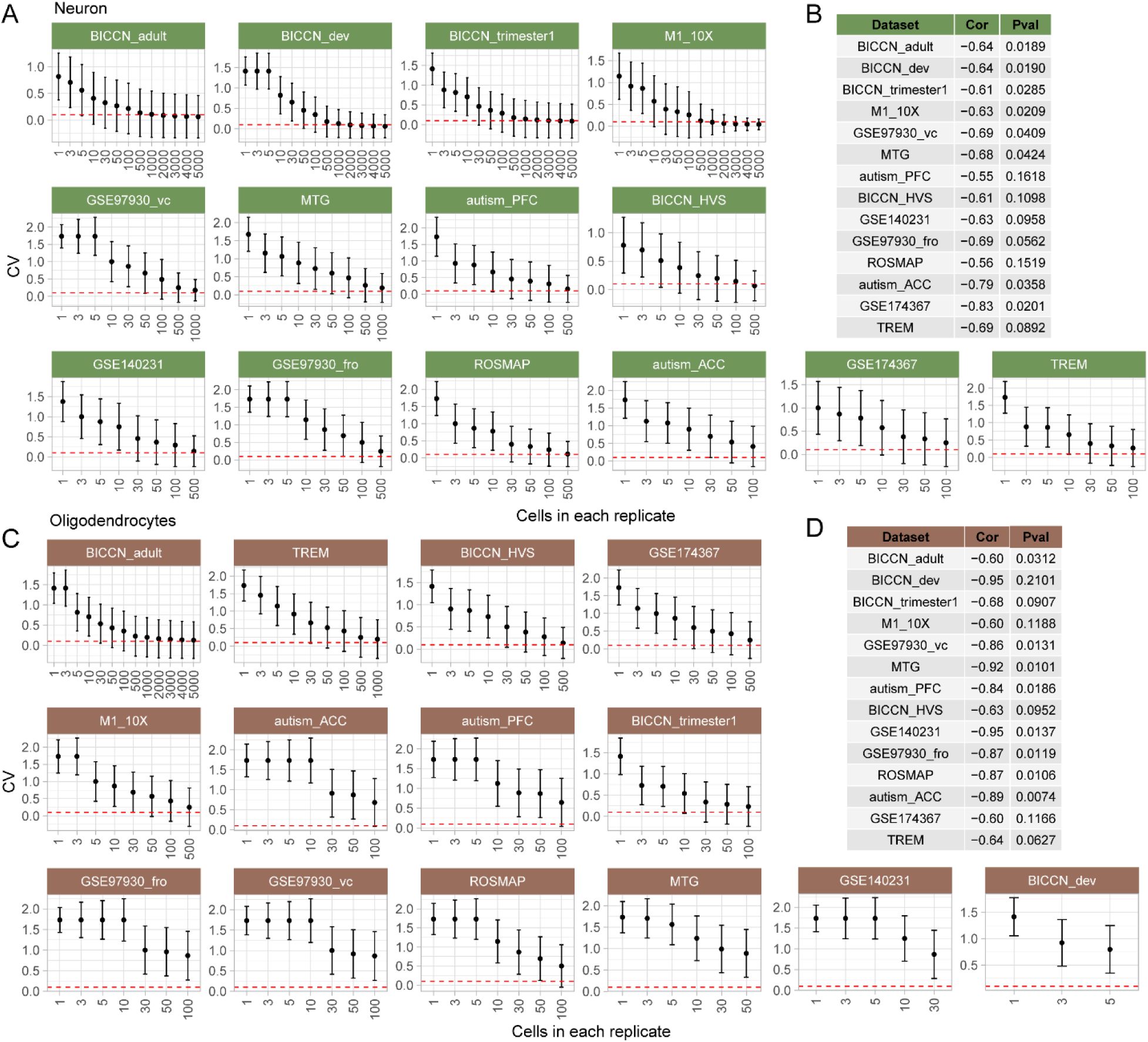
Relationship between cell number and gene expression precision. **A**. CV values in down-sampled neurons. **B**. Pearson correlation coefficient and p value between neuron cell numbers in replicate and median CV. **C**. CV values in down-sampled oligodendrocytes. **D**. Pearson correlation coefficient and p-value between oligodendrocyte cell numbers in replicate and median CV. To enhance visual clarity, the number of cells in each replicate was capped at 5000. The Red dashed line denotes a CV of 0.1. Cor, Pearson correlation coefficient; Pval, p-value.

The minimum number of cells required for delivering acceptable precision is suggested by data of excitatory neurons. Based on five datasets (BICCN_adult, BICCN_dev, BICCN_trimester1, M1_10X, and BICCN_HVS), approximately 500 cells were required to achieve a median CV close to 0.1 for neurons (**Figure 4A**). None of the other cell types attained CV values as low as 0.1 and they all had fewer than 500 cells sequenced.

### RNA integrity is correlated with expression precision

The cell numbers required for achieving an acceptable precision level in sc/snRNA-Seq data vary across studies, suggesting expression precision may not be solely dependent on the number of cells. We examined the effects of four technical factors, including RNA integrity, sequencing depth, sequencing saturation, and sequencing platform, on expression precision of excitatory neurons.

We tested the relationship between RNA integrity, as measured by the RNA integrity number (RIN), and median CV in technical replicates. Two datasets with RIN information available for analysis were used. Samples with higher RIN value tended to have lower CV values (**Figure 5A** and **Figure 5B**). By zooming into replicates with 200 cells, negative correlations were observed between RIN and median CV in the ROSMAP (R^2^ = 0.26, p-value = 0.06, **Figure 5C**) and autism_PFC (R^2^ = 0.60, p-value = 0.04, **Figure 5D**) datasets, suggesting that RNA integrity is another factor contributing to expression precision.

**Figure 5.**
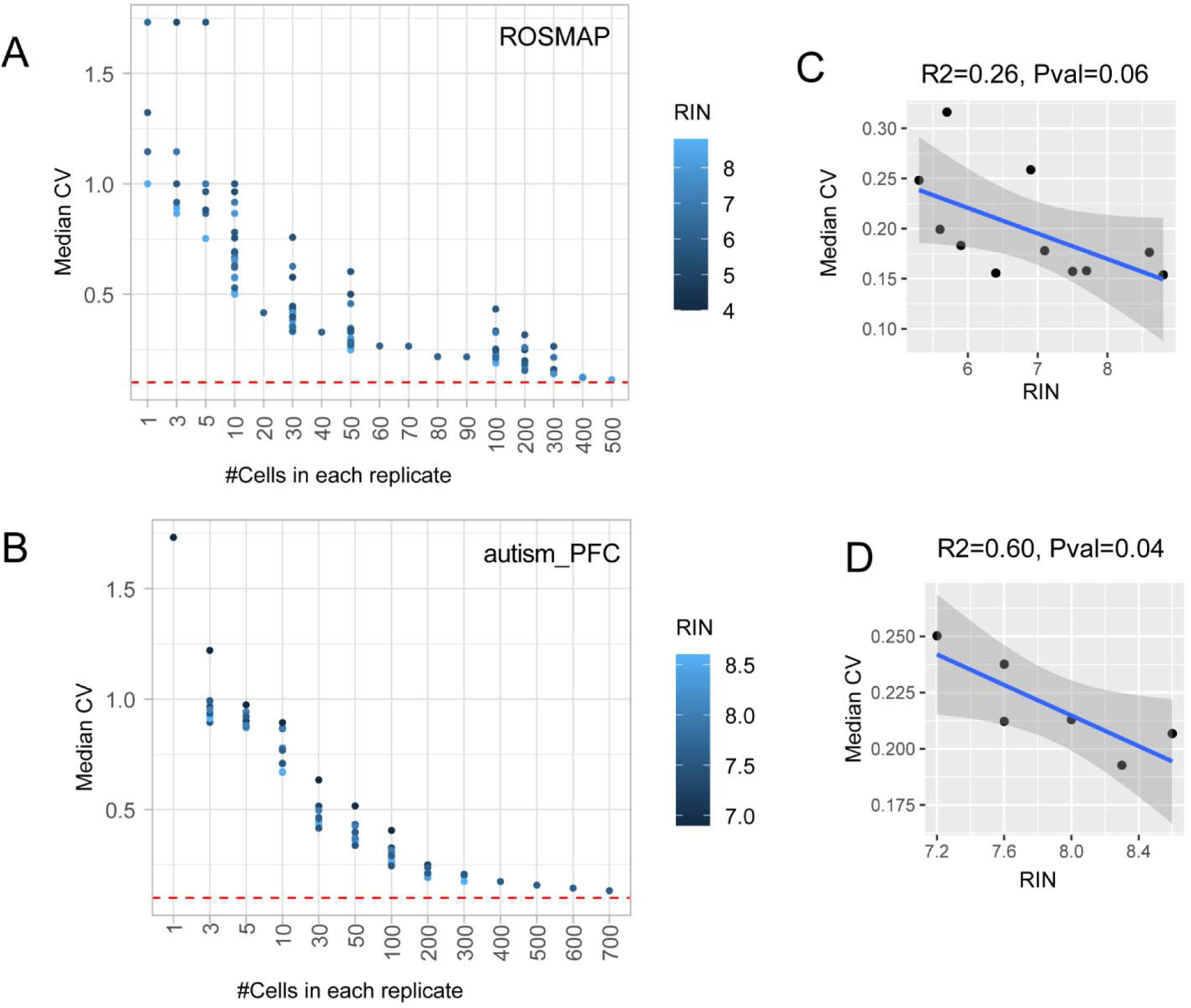
Association between RNA integrity and gene expression variability across technical replicates in snRNA-Seq data. The relationship between number of cells in replicate and median CV was tested in ROSMAP (**A**) and autism_PFC (**B**) dataset. Samples were colored by RIN. Red dashed line denotes the CV threshold of 0.1. (**C**) & (**D**) illustrate the relationship between RIN and median CV when replicate containing 200 cells in ROSMAP and autismPFC data, respectively. Linear regression model was used. RIN, RNA integrity number.

In the autism_PFC data, we also explored the correlation between median CV and total sequencing depth (p-value = 0.89) and saturation rates (p-value = 0.76), but no significant correlation was found (Figure S10A, Figure S10B). We also compared expression variability across technical replicates in data generated from two different sequencing platforms, 10X Chromium (autsim_PFC) and Smart-seq (MTG). The median CV across detected genes in replicates constructed by 200 cells was used for comparison. No significant difference in gene expression variability was observed between the two technologies (p-value = 0.56, Wilcoxon signed-rank test, Figure S10C), indicating the precision problem is not unique to a specific sequencing platform.

### Low expression accuracy in scRNA-Seq data associated with the number of cells sequenced

To evaluate the accuracy of gene expression, we compared pooled-cell RNA-Seq data with single-cell RNA-Seq data of cultured mononuclear phagocytes from matched samples (**Figure 6A**). RNA-seq data from pooled cultured cells (of one type) was referred to as pooled-cell RNA-seq. The gene expressions from pooled-cell RNA-Seq were considered as the ground truth. We used Pearson correlation and linear regression to assess the expression accuracy. In the linear regression model, the ground truth was treated as the independent variable, while the pseudo-bulks from sample-matched scRNA-Seq was the dependent variable. We tested the significance of the slope deviating from one. The significance of the linear regression, combined with the Pearson correlation coefficient, was used to measure expression accuracy. We calculated the expression accuracy independently for each of the four species. To illustrate the relationship between the number of cells and expression accuracy, we performed downsampling experiments, ranging from 1,000 to 1 cell for each sample.

**Figure 6.**
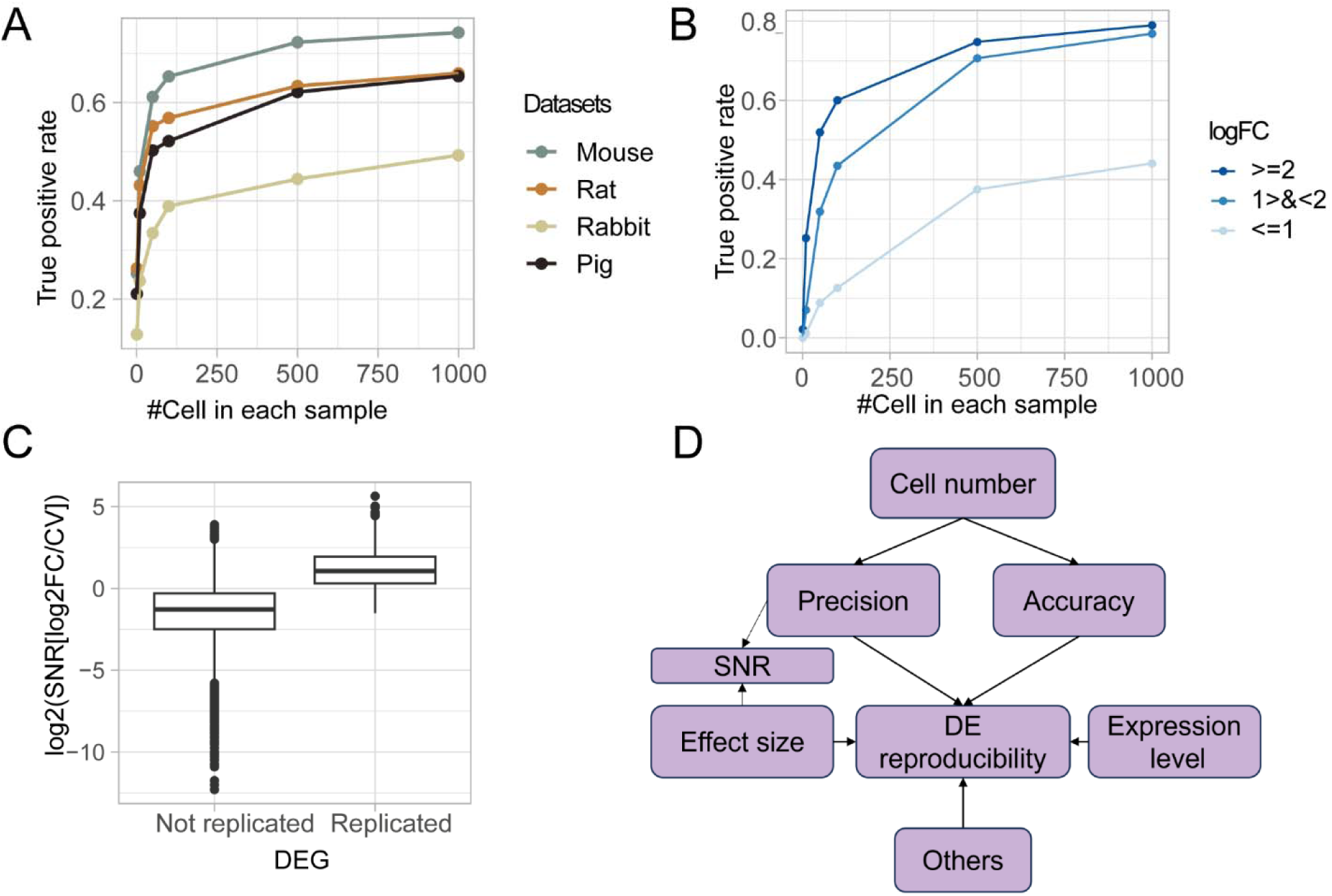
Relationship between expression accuracy and cell number. **A**. Illustration of expression accuracy assessment. **B**. The relationship between cell number and number of genes with good accuracy is defined by Pearson correlation and linear regression model.

The number of genes with good accuracy decreased in down sampling (**Figure 6B**). We observed 3,450 out of 13,907 detected genes with good accuracy as defined by criteria of regression slope of 1 (p-value of 0.05) and correlation coefficient of 0.9 when 1000 cells were analyzed for each sample in mouse data. When each sample contained a single cell, only 100 genes showed good accuracy. When data have 500 cells in each sample, the gene accuracy tends to reach a stable value. Similar patterns were observed in data from rat, pig, and rabbit, though pig and rabbit data showed overall worse performance than mouse and rat data (Table S1).

The relationship between number of cells and expression accuracy was replicated in the simulation data. In the simulation, scRNA-Seq data of six samples, each with 3000 cells, were synthesized. The pseudo-bulks of 3000 cells in each sample were used as ground truth. We observed that the number of genes with good accuracy increased with larger cell numbers (Figure S11), consistent with results from our real data. Notably, when at least 500 cells were sampled, the number of genes with good accuracy began to stabilize.

### Noise level and trait effect size interactively affects the reproducibility of differential expression analysis in scRNA-Seq data

To assess the impact of data quality on downstream analysis, we conducted a DE analysis in sample-matched scRNA-Seq and pooled-cell RNA-Seq datasets independently, comparing Lipopolysaccharide (LPS)-treated and untreated groups using the edgeR algorithm. Genes were considered significantly differentially expressed (DEGs) when their false discovery rate (FDR)-corrected p-value was less than 0.05. Since we already showed that both expression precision and accuracy were positively correlated with the number of cells sequenced, we employed a down-sampling strategy to investigate the influence of cell number on DE results. By utilizing the DE results in pooled-cell RNA-Seq data as ground truth, we evaluated the overall reproducibility of DE results in scRNA-Seq data with true positive rate, the proportion of actual positive instances that are correctly identified as positive. Notably, as the number of cells increased, the true positive rate improved and had a plateau at about 500 cells (**Figure 7A**). The true positive rates were 0.72, 0.63, 0.62, 0.44 in data from mouse, rat, pig and rabbit, when 500 cells were included in each sample.

**Figure 7.**
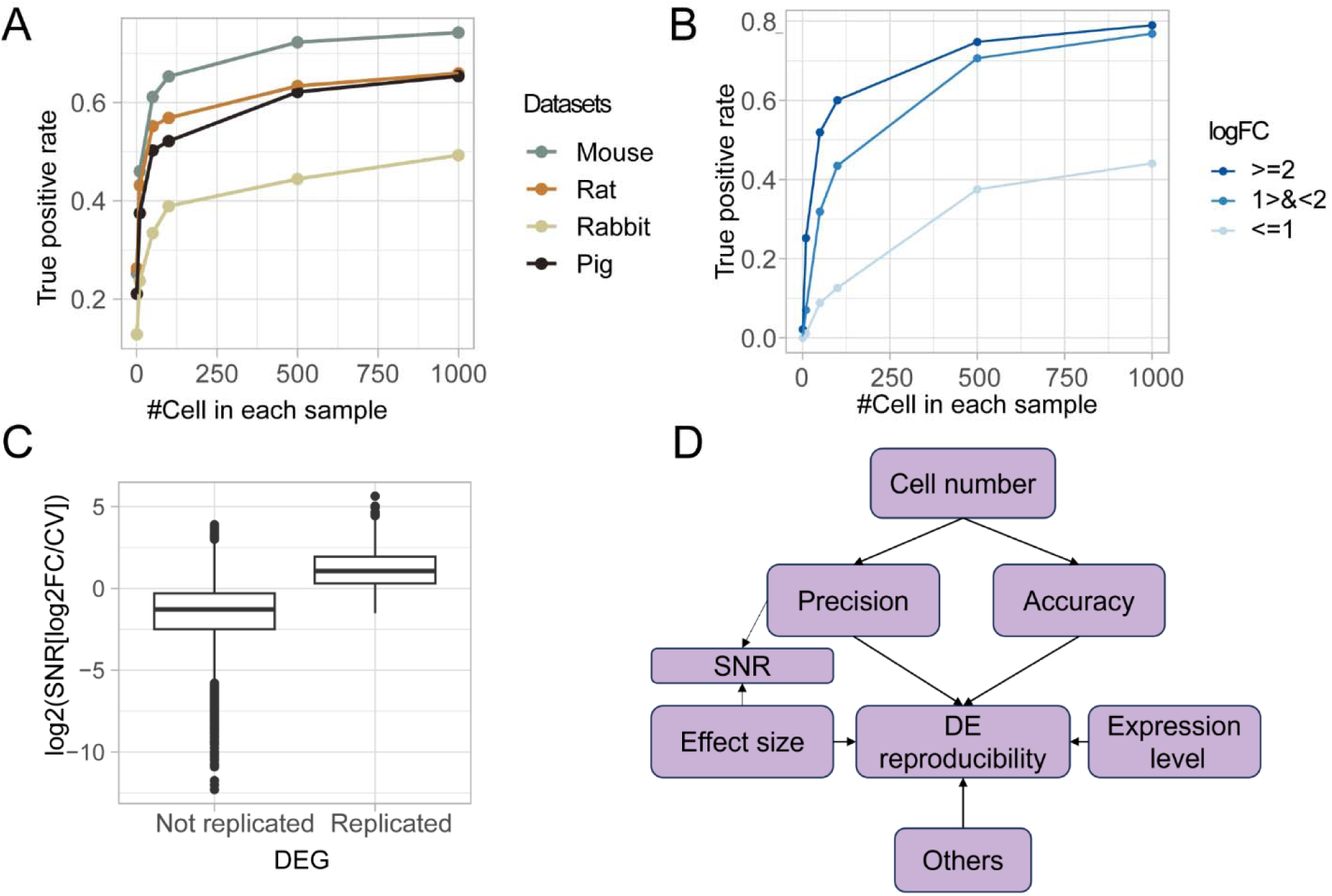
Reproducibility of differential expression analysis in scRNA-Seq data. **A**. True positive rate of DEGs in data with different numbers of cells. Datasets from different species are colored. **B**. True positive rate of DEGs with different effect sizes categorized by logFC of differential expression. **C**. The relationship between signal-to-noise ratio (SNR), and DEG reproducibility in mouse data. **D**. Model of DE reproducibility.

Effect size, which reflects the differences between the compared groups, plays a crucial role in determining the statistical power of a DE analysis. By categorizing the mouse DEGs into three groups based on effect size: high (|log2FC| ≥ 2), medium (1 < |log2FC| < 2), and low (|log2FC| ≤ 1), we observed that DEGs with high and medium effect sizes demonstrate a better true positive rate than those with low effect sizes. DEGs with medium effect sizes still exhibit a relatively lower true positive rate compared to genes with high effect sizes, particularly when the number of cells is limited. (**Figure 7B**).

For example, when 500 cells were included in each sample, DEGs with large effect sizes (over two-fold changes) had a true positive rate of 0.73, whereas DEGs with small effect sizes had a true positive rate of only 0.38. When only 50 cells were included in each sample, the true positive rates were 0.41 and 0.09 for DEGs with large and small effect sizes, respectively.

This suggested that the relationship between effect size and noise level has an interactive impact on DEG reproducibility. To quantify this combined effect, we adopted signal-to-noise ratio (SNR) metric for each gene, defined as normalized effect size divided by CV. Using mouse data as an example, we found that replicated DEGs exhibited significantly higher SNRs than non-replicated DEGs (P < 2.2 × 10□^1^□, **Figure 7C**). This trend was consistent with the observation that DEGs showing higher expressions tend to have better reproducibility (Figure S12). The factors influencing DEG reproducibility are summarized in **Figure 7D**.

To evaluate the applicability of the 500-cell cutoff and the SNR measurement, we applied various cell number cutoffs to an independent dataset from Ruzicka et al.[19], which conducted DE analysis on two schizophrenia postmortem brain cohorts (MCL and Mt Sinai). Using an Exact test, we assessed the reproducibility of DEGs across the two cohorts. At a 500-cell cutoff, cell types with significant DEG reproducibility were clearly distinguishable from those without (Figure S13A). However, reducing the cell-number cutoff to lower thresholds, such as 300 or 100 cells—commonly regarded as acceptable in practice—may result in misleading indications of reproducibility. For instance, the Vip neuron emerged as a potential cell type with replicable DEGs at these lower cutoffs, yet its reproducibility was not statistically significant. Moreover, we found that reproduced DEGs exhibited significantly higher SNR (P = 0.0002, Figure S13B), effectively distinguishing reproducible genes from non-reproducible ones in this dataset.

## Discussion

The use of sc/snRNA-Seq in biological studies has become a common practice, necessitating meticulous evaluation of data quality to avoid misleading or even false findings. The current investigation assesses the expression precision and accuracy of published sc/snRNA-Seq data. By analyzing 23 representative datasets, we demonstrated that the gene expressions per individual measured for most cell types were of low precision and accuracy. We found a robust correlation between the number of cells sequenced and the precision, accuracy, and reproducibility of downstream DE analysis. Only cell types having a large number of cells (minimum 500 cells) sequenced delivered relatively accurate and precise quantification of gene expression and, consequently, credible results of downstream analyses, such as case-control comparisons.

Many studies have speculated that cell number, RNA integrity, and sequencing depth influence sc/snRNA-seq data quality, but none have systematically quantified these effects across datasets or established actionable thresholds. This lack of clear, reproducible standards has led to inconsistencies in experimental design and, in some cases, unreliable—or even outright incorrect—conclusions. A striking example is Murphy et al.[20], who demonstrated that many of the transcriptional differences reported in Mathys et al.[9] regarding Alzheimer’s disease were false positives, caused by inadequate noise control and flawed differential expression analysis.

Alarmingly, this paper —despite its misidentified genes—has been cited over 2,000 times, significantly shaping the Alzheimer’s research landscape. This is just one example; similar issues permeate the field[21–23]. This concern aligns with the findings of previous studies[24, 25]. It is urgent that the single-cell research community recognizes the critical importance of data quality to prevent misleading findings and ensure the reliability of future discoveries. Our study addresses this urgent need by providing a quantitative threshold driven by large datasets, gene-level evaluation metrics, and practical tools and guidance.

Our study establishes quantitative thresholds critical for ensuring high-quality single-cell data and results. Prior studies qualitatively recognized that increasing the number of cells improves data quality and reproducibility[26, 27], but the relationship between them is non-linear and the gene expression precision, accuracy and reproducibility saturate at certain cell numbers (Figure 4A, Figure 6B, and Figure 7A). Therefore, a quantitative cutoff is required to exclude low-quality genes and samples, similar to standard practices in bulk RNA-seq. This cutoff has never been defined, creating a gap that limits consistency and reliability in single-cell studies. Our systematic evaluation of 23 sc/snRNA-seq datasets of matured cell types from brain and other tissues demonstrates that at least 500 cells per cell type per individual are required for robust measurements—an evidence-based threshold previously missing in the field.

The criteria used for evaluating expression precision in this study are standard statistical techniques. We used CV < 0.1 as the cutoff for the expression precision in this evaluation. This is based on previous quality evaluations of bulk RNA-Seq data[16]. We believe that holding sc/sn RNA-seq data to the same standard as bulk RNA seq is appropriate and lowering the standard will lead to noisy results and poor reproducibility. Additionally, our evaluation demonstrated that CV tends to stabilize at around 0.1 in single-cell data, as the number of cells increases. However, when constructing the technical replicates based on pseudo-bulks, we assumed homogeneity within one cell type. Such an assumption can be violated by heterogeneity caused by cell subtypes and states, which may explain the minimum CV that is observed. Nonetheless, our results indicated that cell subtype was not the major cause of poor precision in the cell types we evaluated, as the precision at cell type level is not worse than that at subtype level.

RNA quality is another crucial factor impacting expression precision. Notably, in the BICCN_HVS study[28], the minimum number of cells required to achieve a CV of 0.1 was 500, but other datasets require even larger numbers of cells. A key factor contributing to this difference may be the use of surgical samples in the BICCN_HVS study, as these samples tend to be less degraded than frozen postmortem brain samples. Ensuring high RNA quality in samples, such as using RNA with RIN values greater than 7, will likely reduce the number of cells required for quality quantification.

The signal-to-noise ratio emerges as a pivotal determinant of the reproducibility of DE analysis. Our investigation revealed that DEGs with large effect sizes exhibit superior reproducibility compared to those with smaller effect sizes. When 500 cells were included in each sample where the noise level was low, DEGs with large effect sizes had a true positive rate of 0.73, whereas DEGs with small effect sizes had a true positive rate of only 0.38. When the cell number decreased to 50 cells where the noise level was high, the true positive rates were 0.41 and 0.09 for DEGs with large and small effect sizes, respectively. This comparison indicates that the technical noise matters more for the smaller biological effects and technical noise may be manageable for phenotypes associated with pronounced expression changes. Improving data quality becomes more critical in scenarios where the effect size of the phenotype approaches the noise level. This is particularly relevant for many complex diseases, including neuropsychiatric disorders, where the effect size is typically small[29], necessitating an increase in the number of cells to minimize technical noise.

This work provides gene-level metrics to help refine reliable signals. Most prior studies assessed data quality at the cell or sample level, which can be biased by highly expressed genes. In contrast, our study introduces a gene-by-gene evaluation framework, enabling precise quality and reproducibility assessments for individual genes—a crucial advancement for downstream analyses like differential expression. Specifically, we introduce the SNR as a key metric for assessing DEG reproducibility, calculated by dividing fold change by CV. Applying this approach to schizophrenia DEGs and mouse data, we found that reproduced DEGs have significantly higher SNR, which effectively distinguishes reproducible genes from non-reproducible ones in this data, providing a practical metric for ensuring reliable single-cell data analysis.

We introduce VICE, a powerful tool that enables researchers to assess the quality of existing single-cell data and predict the reliability of differential expression results. With calculated CV values, users can: (1) determine noise levels across different cell types and samples and (2) identify genes with low noise level, ensuring that only high-confidence genes are prioritized for analysis. By inputting cell numbers, effect sizes, and noise levels, VICE estimates the true positive rate for single-cell DE analysis, providing a direct, data-driven framework for optimizing experimental design and result interpretation. For trait-specific analyses, such as DE, VICE can be used to: (1) estimate the true positive rate based on sample size and cell numbers to guide study design; and (2) evaluate DEG reliability by estimating the true positive rate based on signal-to-noise levels rather than relying solely on p-values.

We provide the following guidelines for future single-cell research. For general data quality control, we recommend: (1) prioritizing high-quality RNA samples (RIN ≥ 7) whenever possible, as degraded RNA increases noise and reduces reproducibility; and (2) ensuring sufficient cell numbers for reliable analysis. We suggest at least 500 cells per individual per cell type for optimal precision. If this is not feasible, focusing on genes with low noise levels is advisable. Adjusting the CV threshold based on trait effect size can help balance precision with dataset constraints. For result reporting, we recommend: (1) routinely reporting CV values and associated power as quality metrics in single-cell data analysis; and (2) providing the median CV for each sample in sc/snRNA-seq experiments to assess sample quality. These benchmarks set optimal standards rather than rigid requirements. Researchers can adapt them as needed, using pseudo-bulk strategies for low cell numbers or adjusting CV thresholds based on effect size. Our approach provides practical, data-driven guidance that supports informed decision-making rather than imposing one-size-fits-all rules.

Our findings hold significant implications across multiple domains. Many researchers have reported results from minor cell types in a variety of tissues, but our work casts doubt about the validity of conclusions drawn from much of this research due to insufficient numbers of cells. The accurate identification and comprehensive study of these minor cell types necessitates sequencing more cells. Beyond elucidating the nuances of DE analysis, our results imply low precision and accuracy impacts other analytical methodologies, including cell classification[30], eQTL mapping[31], and the construction of co-expression networks[32]. The effect of cell numbers on other data analyses remains to be explored.

Our study rigorously quantifies these effects and provides concrete, data-driven guidelines to improve sc/snRNA-seq studies. Our goal is not to rescue poor experimental designs—there is no simple fix for flawed data. Instead, we define the scale of the problem with precise numbers, highlighting critical pitfalls in single-cell data analysis. We equip researchers with clear, quantitative metrics to assess which genes and samples meet quality standards for reliable downstream analysis. More importantly, we provide practical tools and data-driven cutoffs, ensuring that future studies are designed correctly from the start, minimizing errors and maximizing reproducibility.

## Conclusion

In this study, we conducted a quantitative evaluation of expression precision and accuracy across 23 representative sc/snRNA-Seq datasets, revealing significant deficiencies in gene expression measurements—particularly when sequencing a limited number of cells. We demonstrate that the reproducibility of DE analysis is tightly correlated with cell number, emphasizing the need for data-driven thresholds in study design. To improve the reliability and reproducibility of sc/snRNA-Seq studies, we recommend sequencing at least 500 cells per cell type per individual, including minor cell types and RNA quality (RIN ≥ 7). Recognizing practical constraints, we provide flexible, evidence-based guidelines rather than rigid requirements. We strongly advocate for quality assessment before downstream analyses to prevent false discoveries. To facilitate this, we developed VICE, a tool that quantifies technical variability, estimates the true positive rate of DE results, and enables data-driven decision-making in sc/snRNA-Seq studies.

## Materials and methods

### Collection of sc/snRNA-Seq data from cortex

A total of 14 brain sc/snRNA-Seq datasets were obtained for analysis. The collected datasets were derived from human brain studies published between 2012 and 2023[8, 9, 28, 34–45]. Samples from individuals with brain disorders were excluded from the analysis to prevent biasing the expression profiles. The raw count and cell annotation data were obtained from the original studies. Due to differences in cell classification across studies, we harmonized cell identities into eight major cell types present in the adult brain, namely excitatory neurons, inhibitory neurons, oligodendrocytes, oligodendrocyte precursor cells, astrocytes, microglia, endothelial cells, and pericytes. The annotation of data from BICCN 2023 collection was retained to evaluate cell subtypes. The scRNA-Seq data of blood, lung, and lymph node from Tabula Sapiens Consortium were used for evaluating expression variability in multiple tissues[17].

### Collection of sample-matched data from four species

To assess expression accuracy, we utilized four sample-matched scRNA-Seq and pooled-cell RNA-Seq datasets. These datasets were sourced from Hagai et al.[46], encompassing bone marrow-derived mononuclear phagocytes derived from mouse, rat, pig, and rabbit, all subjected to stimulation with either lipopolysaccharide or poly-I:C for a duration of four hours. Within each species, a total of three samples received lipopolysaccharide treatment, while three additional samples were designated as control groups. We employed the preprocessed data provided by Squair et al.[24], which is available at https://doi.org/10.5281/zenodo.5048449.

### RIN

The RIN is a numerical value that measures the quality of RNA samples. It is calculated before sequencing using an automated analysis of RNA molecules through electrophoresis, such as Agilent 2100 bioanalyzer. The RIN scale ranges from 1 to 10, with 10 indicating fully intact RNA and 1 indicating completely degraded RNA. We obtained the RIN of samples from the original studies.

### Processing of sc/snRNA-Seq data

The sc/snRNA-Seq data underwent processing using Seurat version 4[47]. The raw count matrix and cell annotation matrix were used as input to Seurat. We filtered out genes with zero expression in more than 1/1000 of the total cells in each dataset. The proportion of transcripts mapped to mitochondrial genes was calculated for each cell, and cells with 10% or more mitochondrial gene expression were removed to prevent the inclusion of dead cells. Additionally, cells with less than 200 detected genes or those with more than three standard deviations from the mean number of detected genes were excluded. The count data were normalized based on library size and were scaled with a factor of 10,000. The normalized data were then log-transformed.

### Marker gene identification

To identify marker genes in the ROSMAP dataset, we utilized a one versus second high strategy at both the cell and pseudo-bulk level. At the cell level, marker genes were identified using Seurat. Genes with a proportion of zero expression greater than 15% in the target cell type were removed prior to marker gene identification. The Wilcoxon signed-rank test was used to assess the expression difference, and genes with a log2FC greater than 1 and FDR-corrected p-value less than 0.05 were defined as marker genes. At the pseudo-bulk level, pseudo-bulks were constructed by aggregating gene expression for the same cell type from the same individual. Marker genes were then tested using DESeq2[48], and the likelihood ratio test was utilized to evaluate the expression difference between the two cell groups. Marker genes with a log2FC greater than 2 and FDR-corrected p-value less than 0.05 were defined as marker genes at the pseudo-bulk level. Finally, marker genes supported by both cell- and pseudo-bulk-level tests were selected as final marker genes.

### Technical replicate construction and CV calculation

To generate technical replicates based on pseudo-bulks for each cell type, cells in the count matrix were randomly divided into three groups for the same individual. The count expression for each gene was then summed within each group. The CV value was calculated for each gene using the following formula:

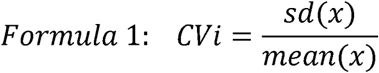

where x represents the gene expression of gene i across three replicates of a specific cell type.

To ensure the robustness of technical replicates, the cell groupings and CV calculations were repeated 100 times, and the average CV across 100 samplings was used.

### CV-cell-number relationship in data with cell class and subclass annotations

To compare the relationship between CV values and the number of sequenced cells in the mouse class and subclass data, a Student’s t-test was used (Figure S3). To compare the relationship between CV values and the number of sequenced cells in the mouse class and subclass data when different number of cells were sequenced, we used a two-sample Z-test (Figure S4). The null hypothesis was that the slope in the regression model testing the relationship between the number of cells and CV values was the same in the class and subclass data. We calculated the difference in slope between the two datasets using the following formula:

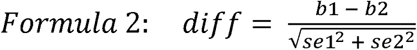

where b1 and b2 were the coefficients, and se1 and se2 were the standard errors from the regression model in the class and subclass data, respectively. We then used the area of the standard normal curve corresponding to the calculated difference to determine the probability in a two-tailed manner.

### Data simulation

Single-cell count data was simulated based on a negative binomial model using the R package Splatter[49]. Two conditions were generated with the “group” simulation, with between 10 and 3000 cells per sample and three replicates per condition. The proportion of differentially expressed genes (‘de.prob’) was set to 0.25.

### Evaluation of expression accuracy

Since no single statistic is sufficient to describe accuracy, we developed a composite criterion that captures the bias and distance from ground truth simultaneously using Pearson correlation and a linear regression model. The scRNA-Seq data were summarized by pseudo-bulks first. The pseudo-bulks were normalized by the library size and were transformed into log2-transformed counts per million (CPM). Then the batch effect between pseudo-bulks and pooled-cell RNA-Seq data was corrected using the combat function in the sva package[50]. Pearson correlation between sample-matched scRNA-Seq and pooled-cell RNA-Seq was calculated for each gene. In the linear regression model, the expression in pooled-cell RNA-Seq was treated as the independent variable and expression in scRNA-Seq was treated as the dependent variable. The intercept of linear regression model was set to 0. By setting offset in function lm in R, the significance of slope deviating from 1 was tested. Good accuracy was defined as correlation coefficient over 0.9 and p-value of linear regression over 0.05.

### DE analysis

DE analysis was carried out on both scRNA-Seq and pooled-cell RNA-Seq datasets to examine the expression disparities between samples treated with lipopolysaccharide and the control samples. For scRNA-Seq data, DE analysis was performed on pseudo-bulk data using the likelihood ratio test approach provided by edgeR[51]. For pooled-cell RNA-Seq data, edgeR was performed on the count data directly. Genes exhibiting an FDR corrected p-value of less than 0.05 were classified as DEGs. We assessed the consistency between DE results obtained from single-cell and pooled-cell RNA-Seq with true positive rate which denotes the proportion of DEGs identified in pooled-cell RNA-Seq that were also replicated in the scRNA-Seq data.

### Application of DE analysis to Ruzicka et al

The schizophrenia DE results were obtained from the supplementary materials of Ruzicka et al. This study includes samples from the Mt Sinai and MCL cohorts. DEGs were defined as genes with log FC > 0.1 and FDR < 0.05. Replicated DEGs were those that met the DE criteria in both cohorts. The Exact test[52] was performed to assess the statistical significance of DEG replication.

## Data and code availability

The data supporting the findings of this study are publicly available, with details for accessing the datasets provided in **Table 1**. The code for this paper can be found at https://github.com/RujiaDai/VICE and https://ngdc.cncb.ac.cn/biocode/tool/BT7673.

## Declaration of AI and AI-assisted technologies

During the preparation of this work the authors used ChatGPT to improve the clarity, grammar, and readability of the text. After using this tool/service, the authors reviewed and edited the content as needed and take full responsibility for the content of the publication.

## CRediT author statement

Rujia Dai, Chunyu Liu, and Chao Chen contributed to conceptualization. Rujia Dai, Tianyao Chu, Ming Zhang contributed to formal analysis, data curation, validation, software and methodology. Rujia Dai and Chunyu Liu contributed to the writing of the original draft. Chunling Zhang, Richard Kopp, Kefu Liu, Xusheng Wang, Yue Wang contributed to the investigation, review and editing. Chunyu Liu, and Chao Chen contributed to funding acquisition, supervision, and project administration.

## Acknowledgment

We thank all the participants in studies for making the data available. We also thank Sidney Wang for invaluable guidance and expertise in the experimental design and data analysis. This work was supported by NIH grants U01MH122591, U01MH116489, R01MH110920, U01MH103340, and R01MH126459; the National Natural Science Foundation of China 82022024 and 31970572; the science and technology innovation Program of Hunan Province 2021RC4018 and 2021RC5027. This work was supported in part by the Bioinformatics Center, Xiangya Hospital, and Central South University.

## Supplementary materials

**Figure S1.**
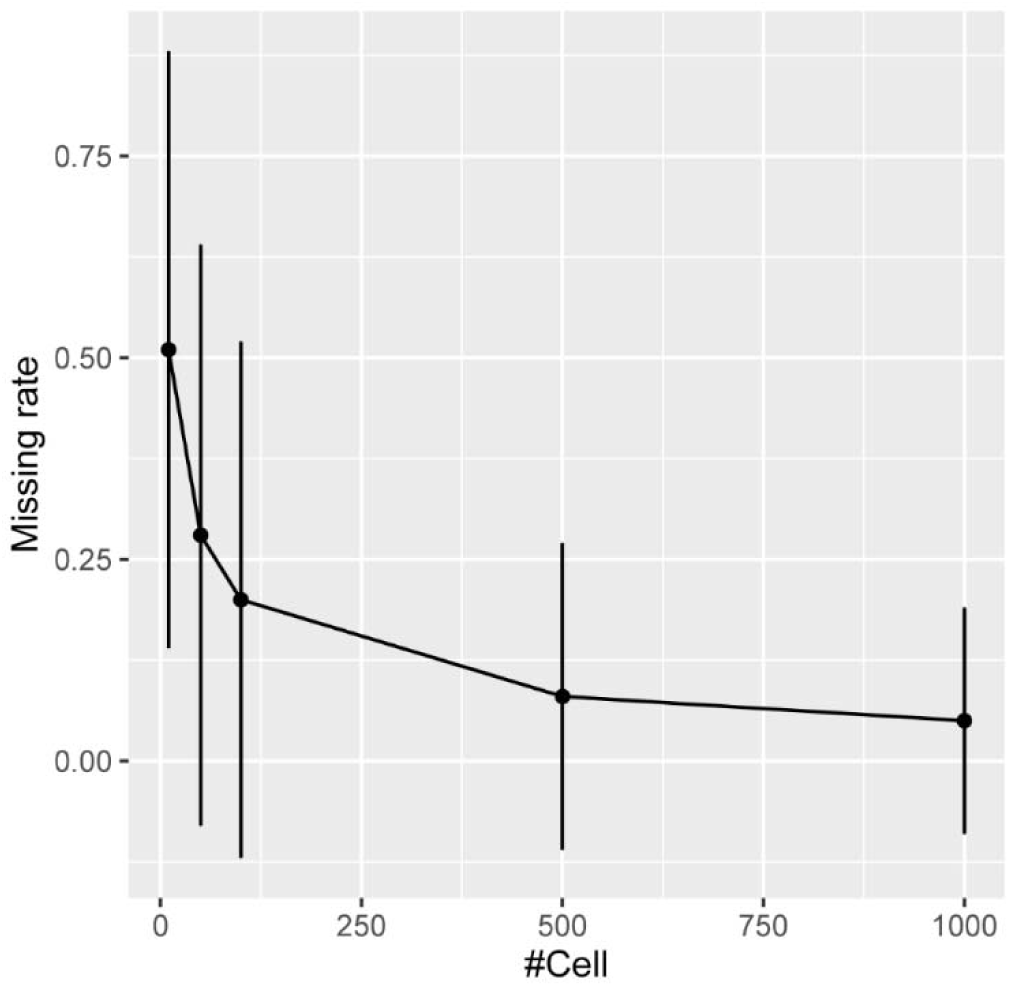
Per gene missing rate in pseudo-bulk data with different number of cells. Missing rate was defined as proportion of zeros in all samples. Data from ROSMAP was used for the demonstration.

**Figure S2.**
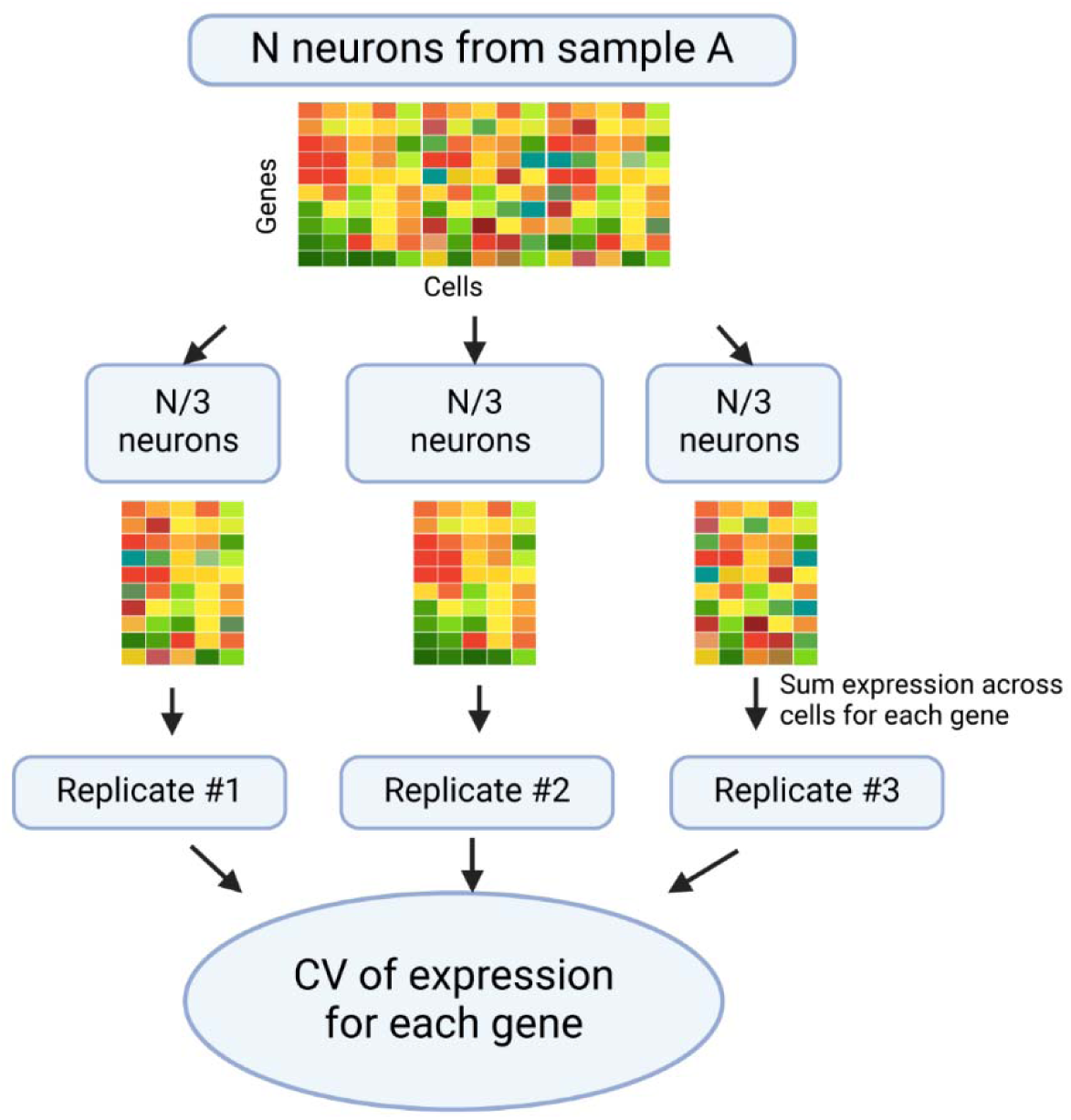
Illustration of constructing technical replicates and calculating the CV of expression across technical replicates. CV, coefficient of variation.

**Figure S3.**
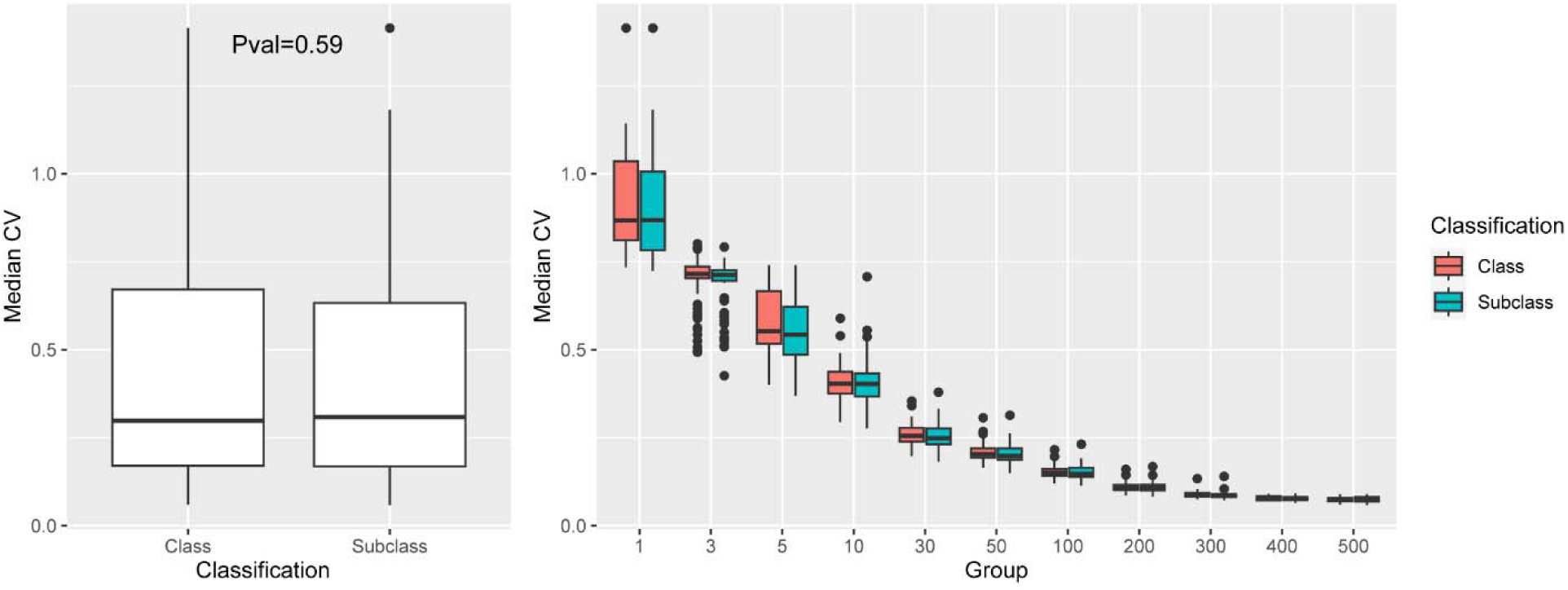
Expression variability at class and subclass level for excitatory neurons. Data from BICCN_HVS was used. P value was calculated with Student’s T-Test.

**Figure S4.**
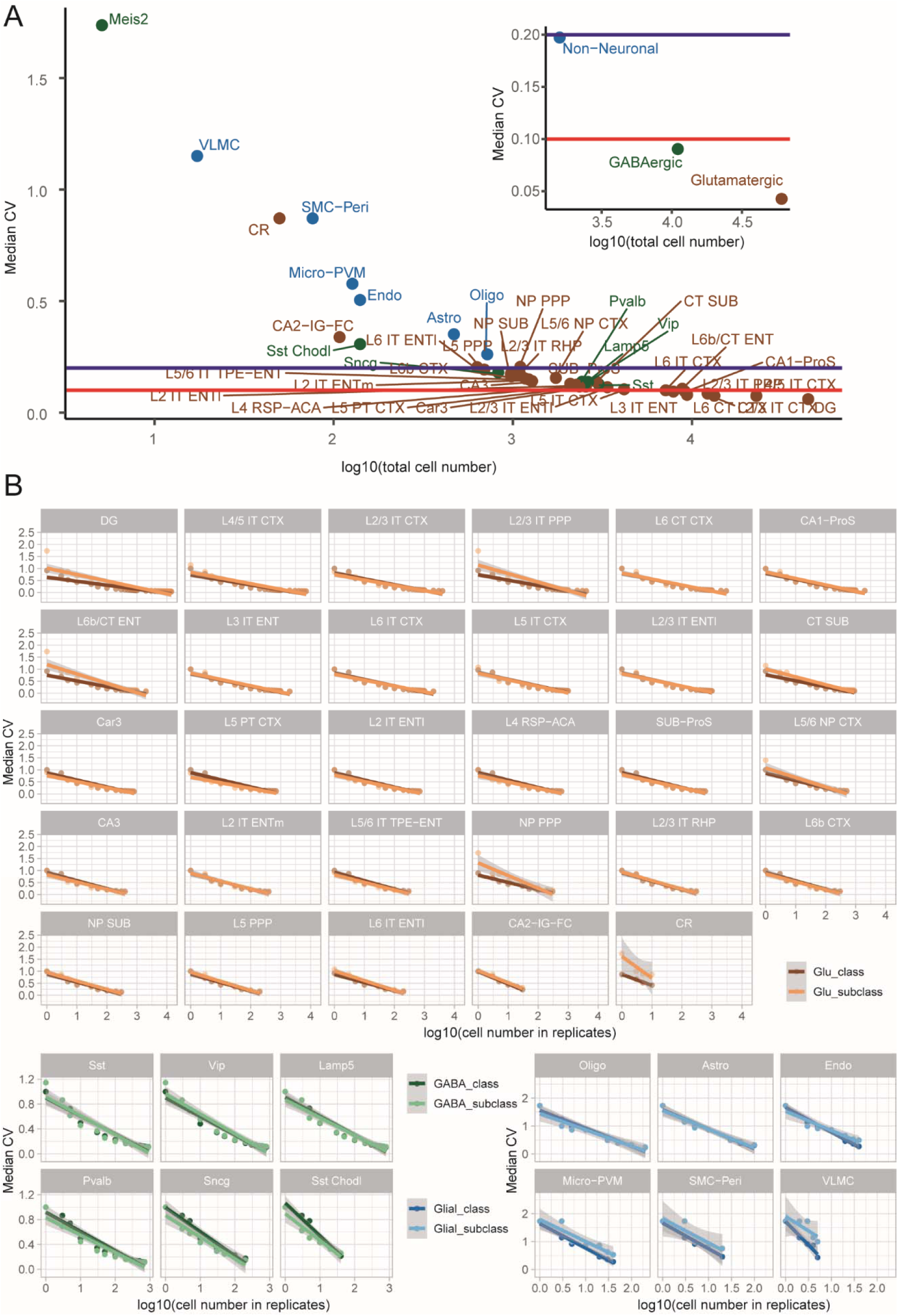
Gene expression variability across technical replicates in mouse brain snRNA-Seq data. **A**. Median CV across genes in replicates constructed at class and subclass level. **B**. Linear regression model of cell numbers in replicates and median CV identified at class and subclass level.

**Figure S5.**
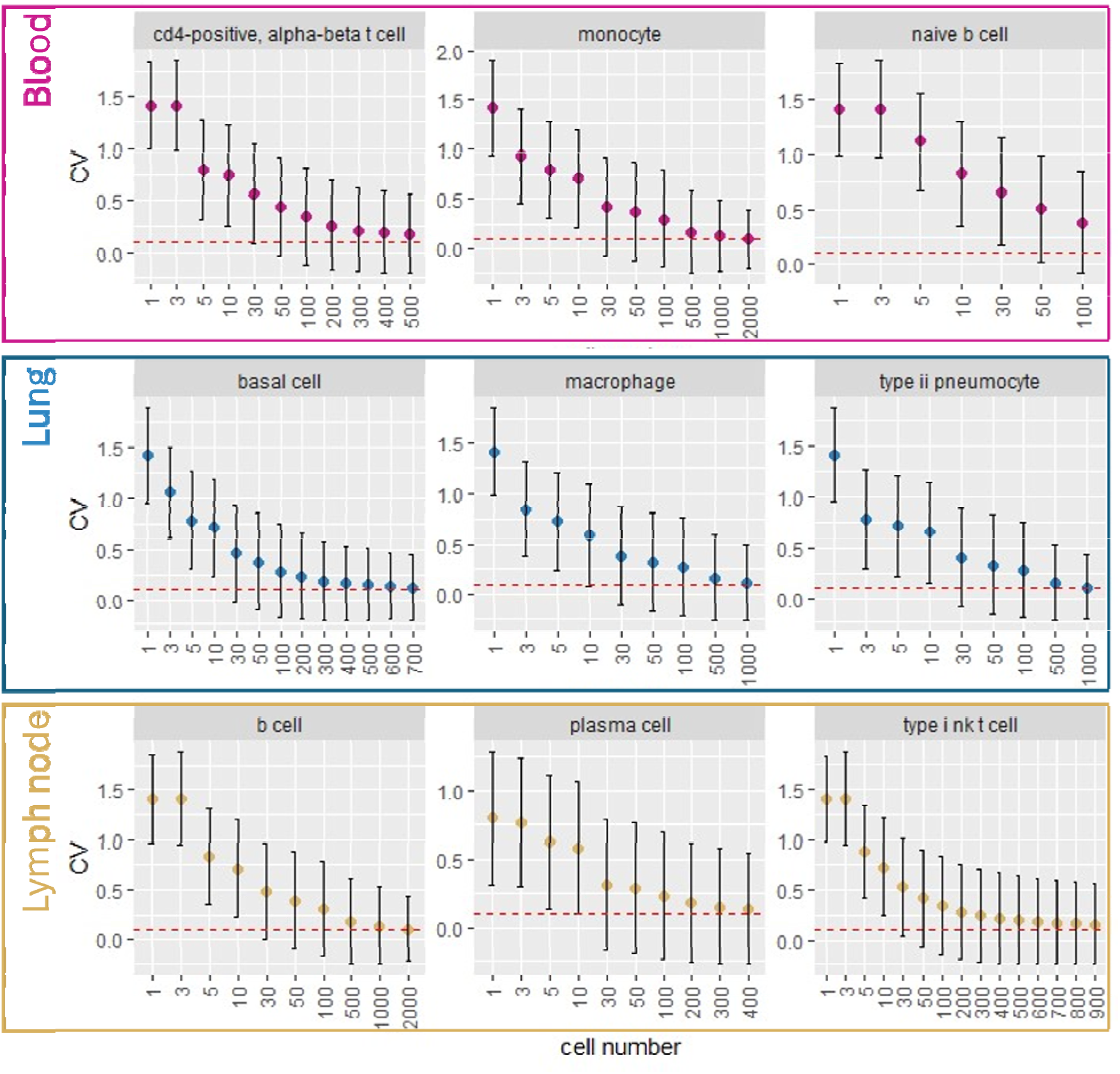
Reduction of CV with the increased number of cells sequenced in three non-brain tissues. Data from Tabula Sapiens Consortium was used. Red line denotes CV of 0.1.

**Figure S6.**
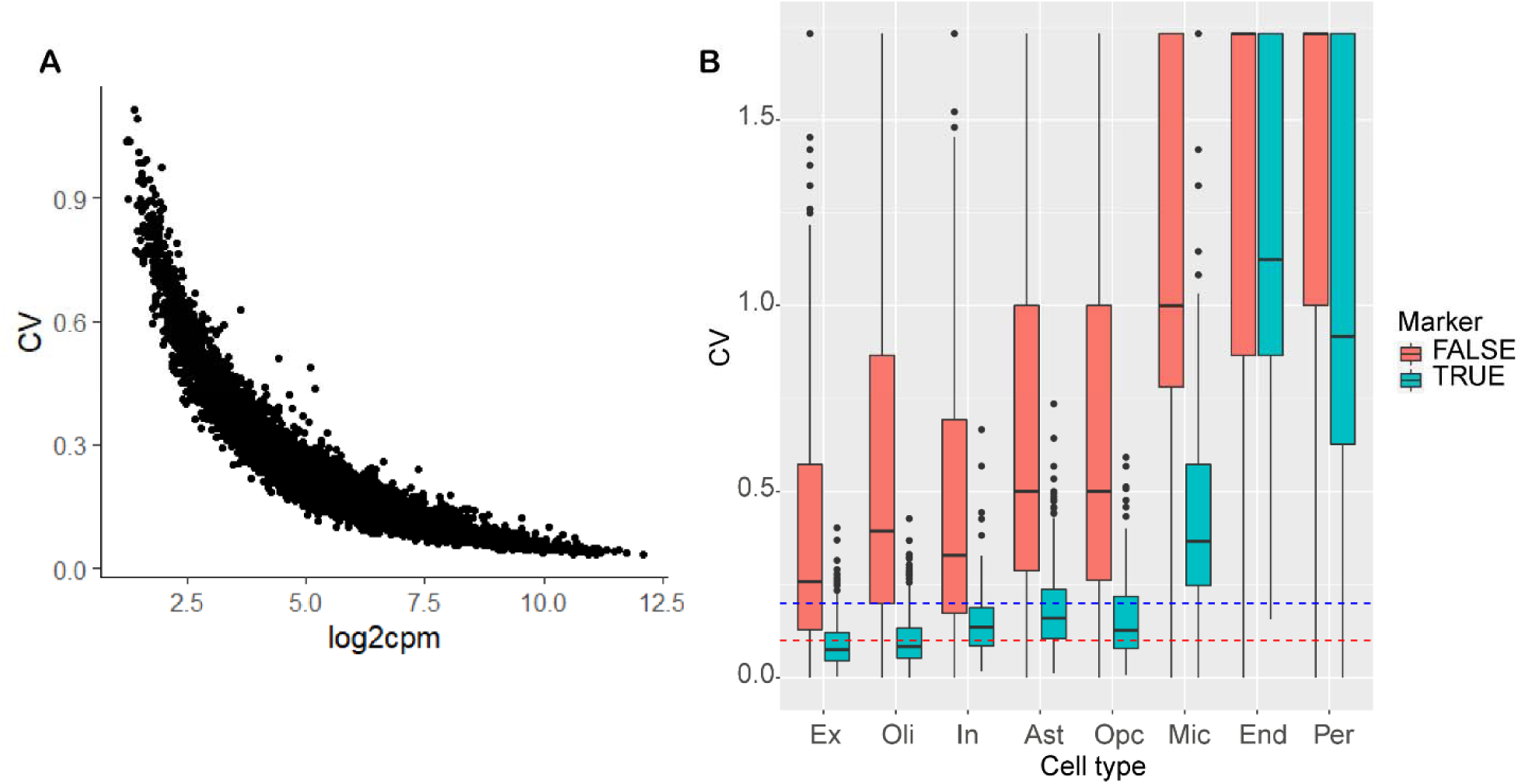
Expression abundance and CV in ROSMAP data. **A**. Relationship of expression and CV in excitatory neuron from ROSMAP study. **B**. CV of marker genes. Blue and red line denote CV of 0.2 and 0.1.

**Figure S7.**
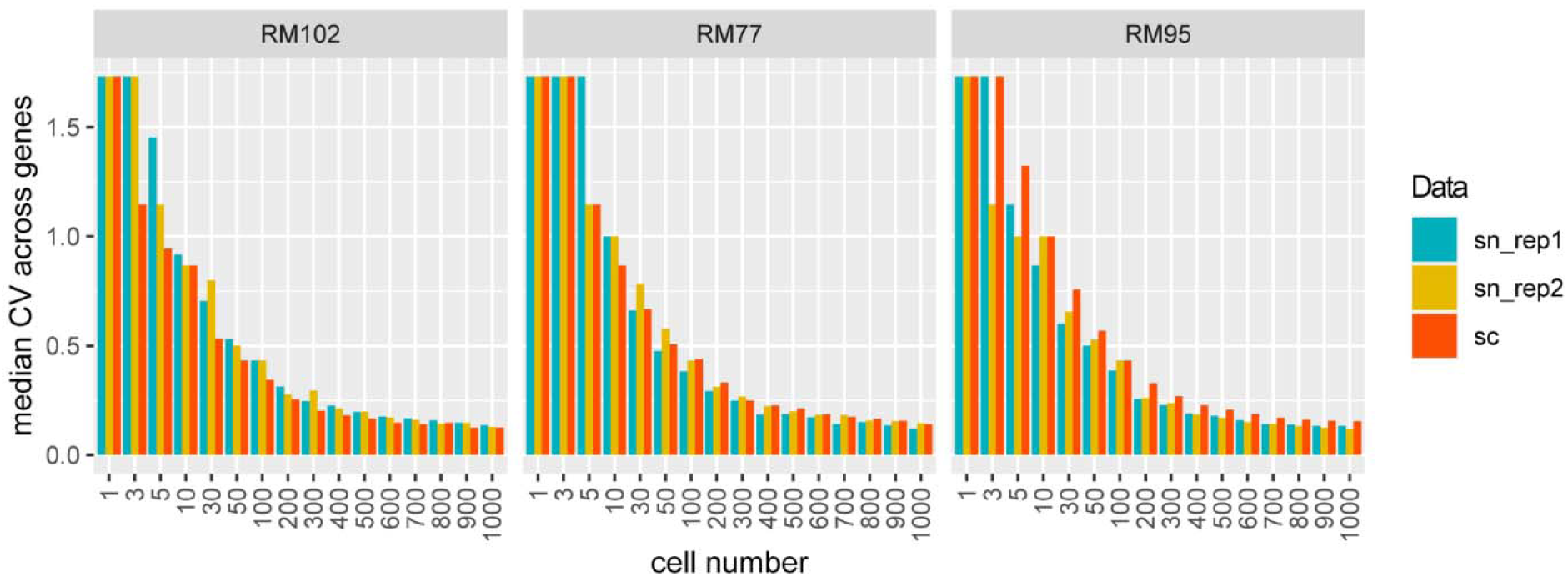
Expression variability in scRNA-seq and snRNA-seq data. Three human microglia samples with both sc- and snRNA-Seq data are shown.

**Figure S8.**
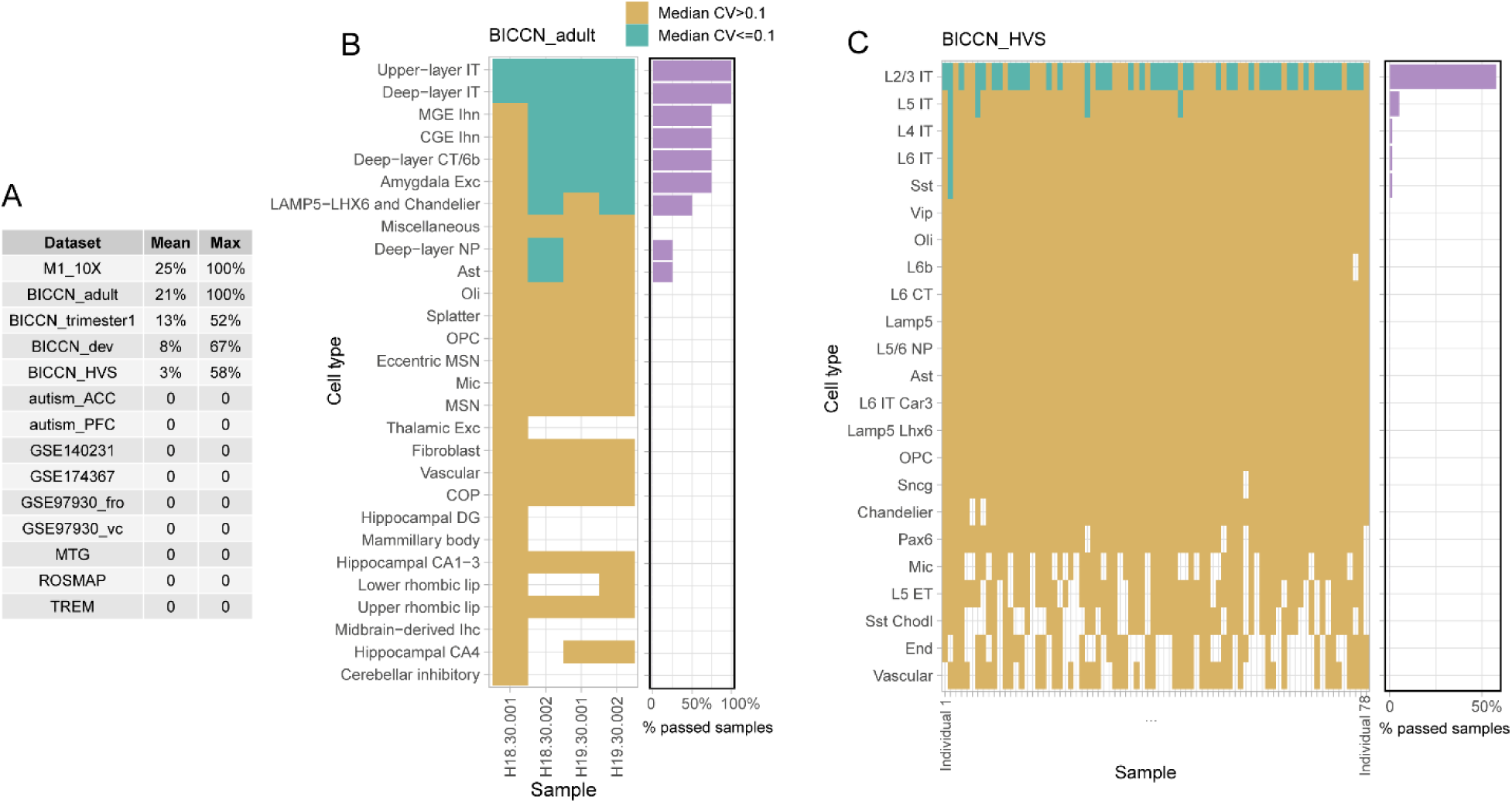
Proportions of samples with acceptable expression variability. **A**. The average and maximum percentages of samples that satisfy the precision criterion. Examples from BICCN_adult (**B**) and BICCN_HVS(**C**) datasets were used for illustration. Samples achieving the precision threshold, defined by a CV of 0.1 or lower, are indicated in green, signifying acceptable expression precision, while those failing to meet the threshold are marked in yellow, indicating low precision. Instances where a cell type is not represented in a sample are left blank. The accompanying bar plot provides a detailed breakdown of the exact proportion of samples that satisfy the precision criterion.

**Figure S9.**
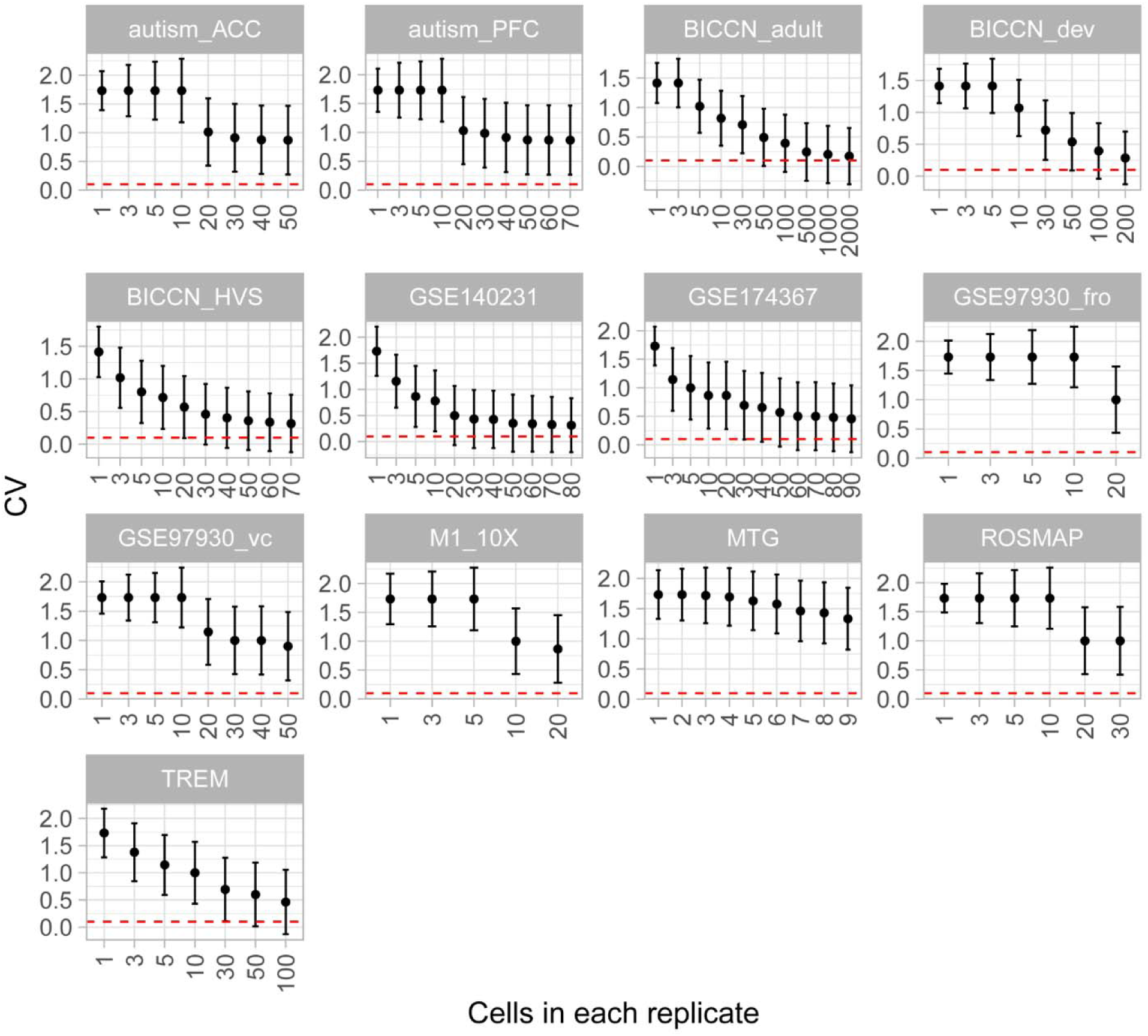
Expression variability across replicates in human microglia.

**Figure S10.**
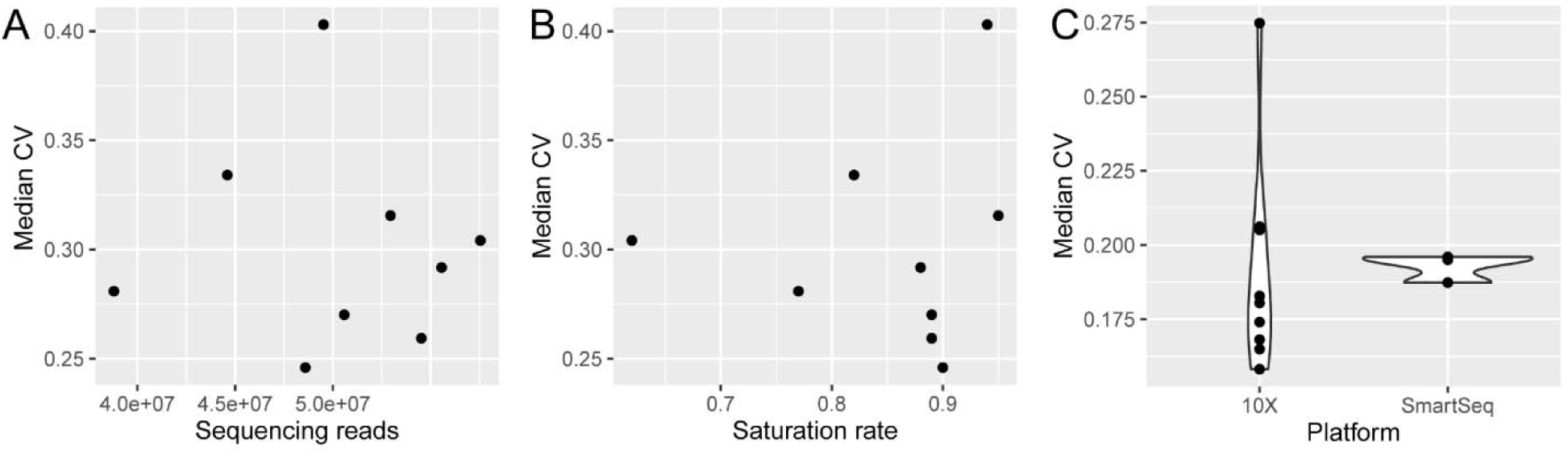
Association between technical factors and gene expression variability across technical replicates in snRNA-Seq data. **A**. Median CV across detected genes in replicates versus total sequencing depth. **B**. Median CV across detected genes in replicates versus total sequencing saturation rate. **C**. Comparison of median CV across detected genes in replicates between data sequenced by 10X Chromium and Smart-Seq platforms.

**Figure S11.**
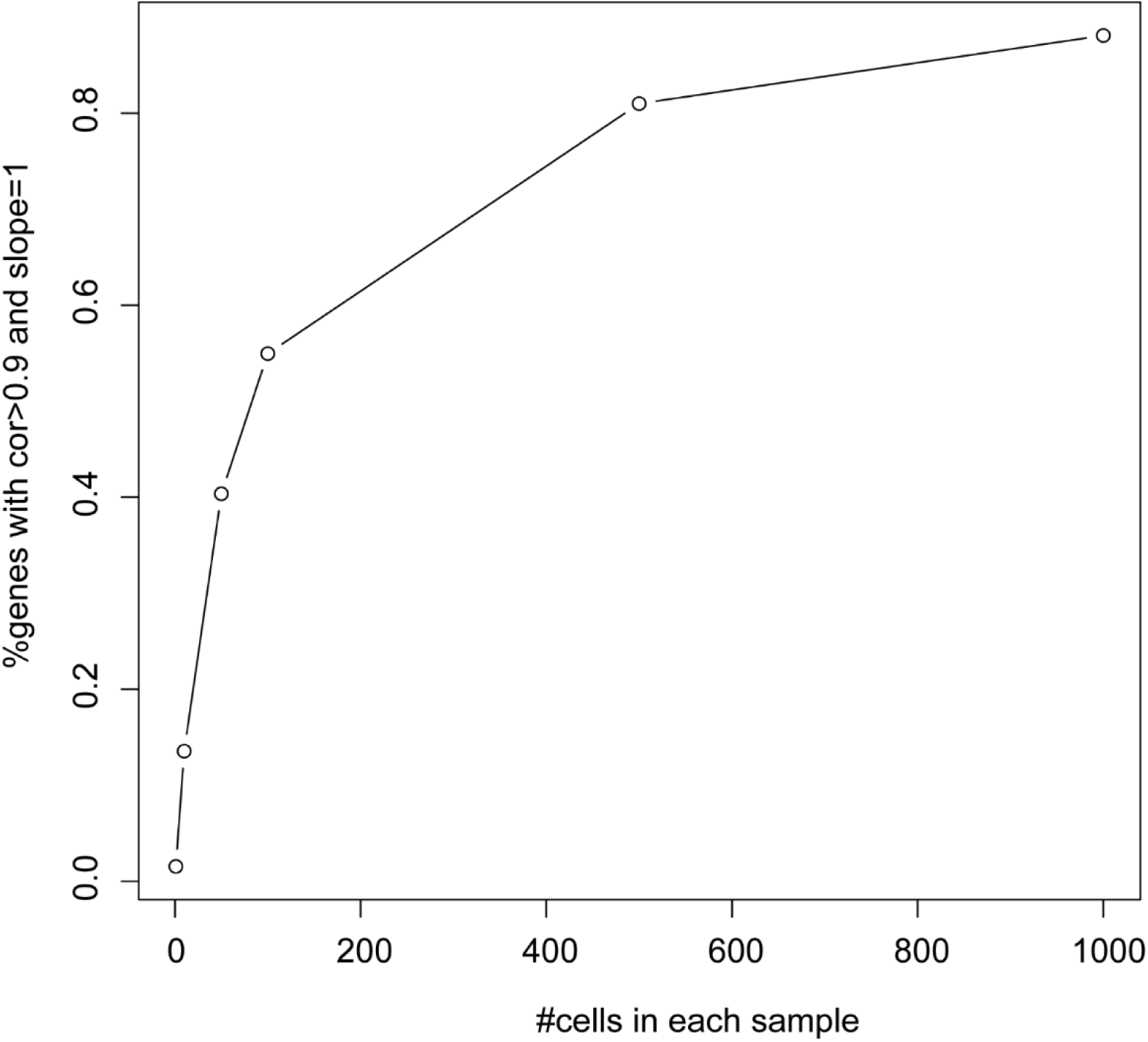
Relationship between expression accuracy and the number of cells in simulated data. The X-axis represents the number of cells in each sample, and the Y-axis shows the percentage of genes with good accuracy, as defined by Pearson correlation and a linear regression model.

**Figure S12.**
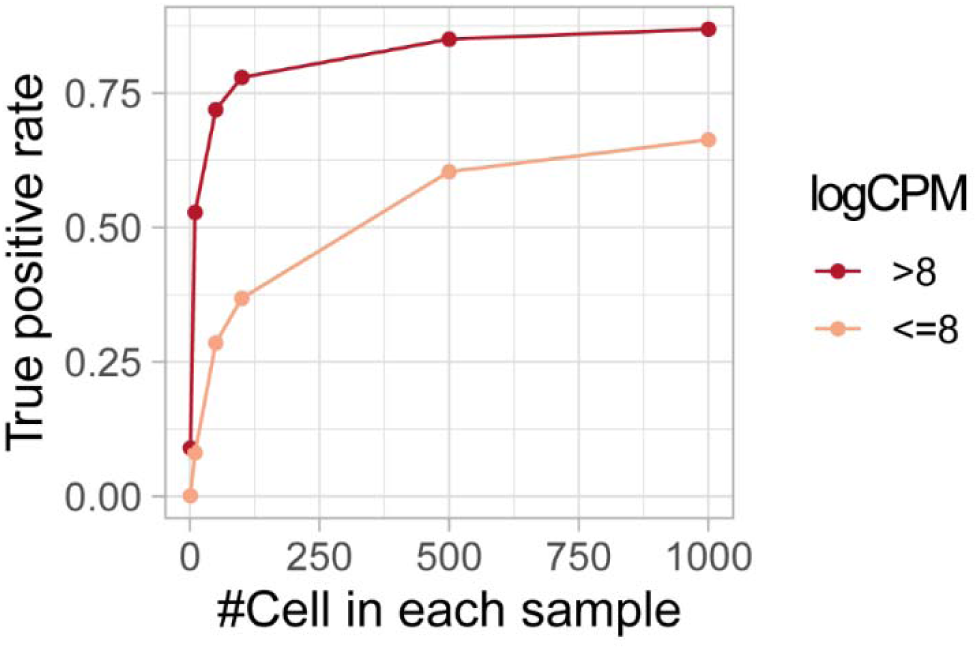
True positive rate of DEGs with different expression levels categorized by log-transformed counts per million (CPM).

**Figure S13.**
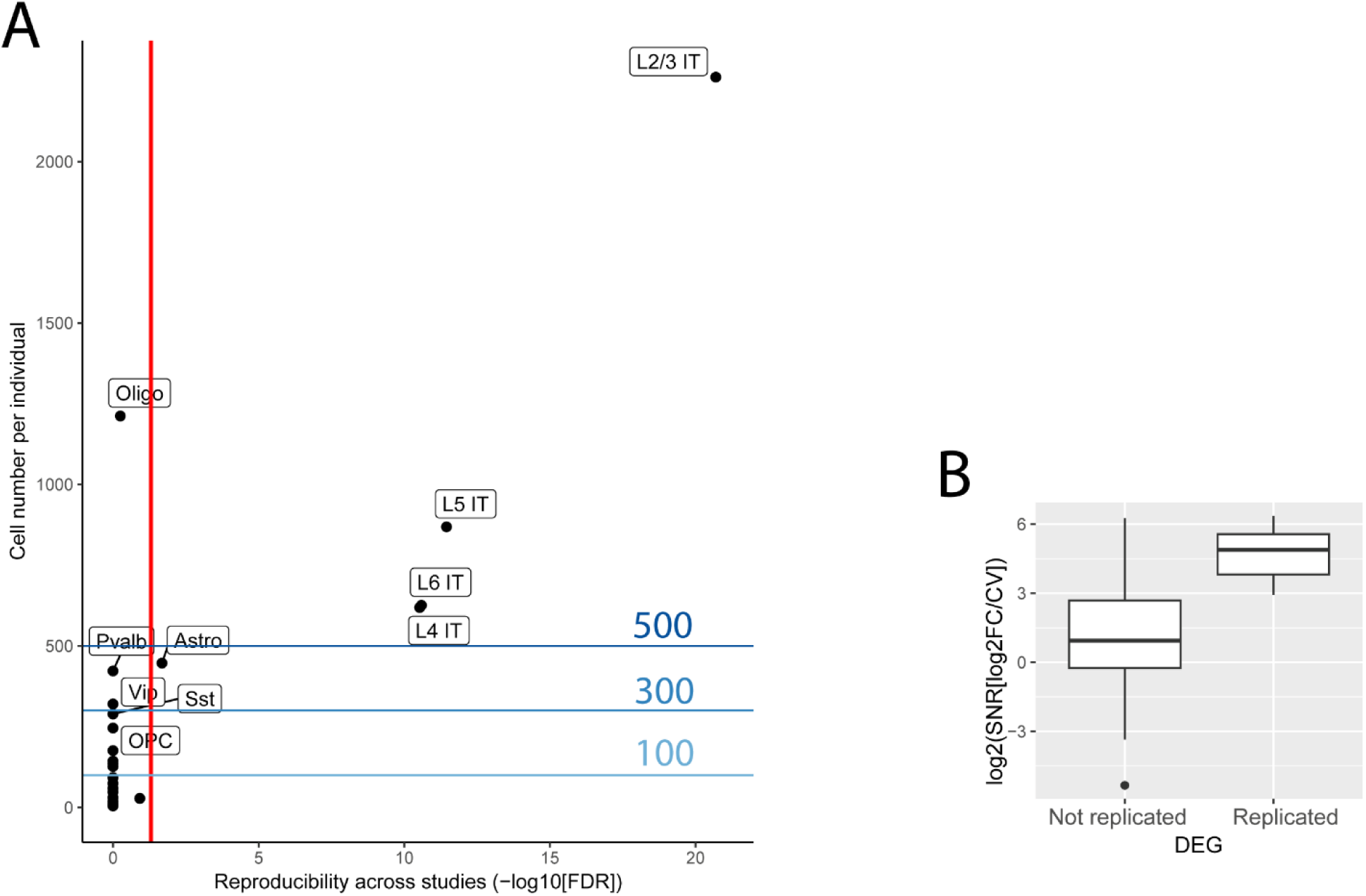
Applying the 500-cell threshold and SNR to schizophrenia case-control scRNA-seq data (Ruzicka et al). **A.** Impact of cell number cutoff on the reproducibility of DEGs in two schizophrenia cohorts. The plot illustrates the effect of different cell number cutoffs on the reproducibility of DEGs identified in two independent schizophrenia cohorts (MCL and Mt Sinai). **B.** The relationship between SNR and DEG reproducibility in astrocytes.

**Table S1 Expression accuracy statistics in datasets from four species**

